# Identifying mutations in *sd1*, *Pi54* and *Pi-ta,* and positively selected genes of TN1, the first semidwarf rice in Green Revolution

**DOI:** 10.1101/2021.12.23.474023

**Authors:** Jerome P. Panibe, Long Wang, Yi-Chen Lee, Chang-Sheng Wang, Wen-Hsiung Li

## Abstract

**Background:** Taichung Native 1 (TN1) is the first semidwarf rice cultivar that initiated the Green Revolution. As TN1 is a direct descendant of the Dee-geo-woo-gen cultivar, the source of the *sd1* semidwarf gene, the *sd1* gene can be defined through TN1. Also, TN1 is susceptible to the blast disease and is described as being drought-tolerant. However, genes related to these characteristics of TN1 are unknown. Our aim was to identify and characterize TN1 genes related to these traits.

**Results:** Aligning the *sd1* of TN1 to Nipponbare *sd1*, we found a 382-bp deletion including a frameshift mutation. Sanger sequencing validated this deleted region in *sd1*, and we proposed a model of the *sd1* gene that corrects errors in the literature. We also predicted the blast disease resistant (R) genes of TN1. Orthologues of the R genes in Tetep, a well-known resistant cultivar that is commonly used as a donor for breeding new blast resistant cultivars, were then sought in TN1, and if they were present, we looked for mutations. The absence of *Pi54*, a well-known R gene, in TN1 partially explains why TN1 is more susceptible to blast than Tetep. We also scanned the TN1 genome using the PosiGene software and identified 11 genes deemed to have undergone positive selection. Some of them are associated with drought-resistance and stress response.

**Conclusions:** We have redefined the deletion of the *sd1* gene in TN1, a direct descendant of the Dee-geo-woo-gen cultivar, and have corrected some literature errors. Moreover, we have identified blast resistant genes and positively selected genes, including genes that characterize TN1’s blast susceptibility and abiotic stress response. These new findings increase the potential of using TN1 to breed new rice cultivars.

## Background

The Green Revolution (GR) in rice production was attributed to the high-yielding semi-dwarf cultivars. In fact, the miracle rice, IR8, inherited the *sd1* (semidwarf 1) gene from the Dee-geo-woo-gen (DGWG) cultivar (Hargrove et al. 1979). It conferred IR8 its short stature, making it lodging resistant, leading to high grain yield. Unknown to many, another cultivar also inherited the *sd1* gene directly from DGWG. It is the Taichung Native 1 (TN1), which was popular in the 1960s (Chandler 1992). Recently, the genome of TN1 was sequenced, assembled and annotated, helping to answer questions about the yield difference between TN1 and IR8 and why they both are photoperiod-insensitive (Panibe et al. 2021).

A fundamental characteristic of TN1 is its short height due to the *sd1* gene from DGWG. The deletion of the semidwarf *sd1* gene incurs a loss of function for the gibberellin (GA) 20-oxidase 2 (Os20ox2), which is involved in the synthesis of the growth hormone gibberellin (Spielmeyer et al. 2002). A reduction in GA results in a shorter plant height (Itoh et al. 2002). However, the sequence of the *sd1* gene is not well studied. The current literature definition of the *sd1* gene was based on the comparison of DGWG-type *sd1* mutants (Habataki, Milyang 23, and IR24) with the *sd1* of Nipponbare, Sasanishiki, and Calrose (Monna et al. 2002). It revealed a 383-bp deletion from the second half of Nipponbare’s exon 1 to the first half of exon 2, or in terms of the expressed sequence, a 278-bp deletion (Monna et al. 2002). Another definition of the *sd1* deletion is a 280-bp deletion in the comparison of the semidwarf Doongara with the tall Kyeema, whose *sd1* sequence is similar to Nipponbare (Spielmeyer et al. 2002). Those studies were done when the full Nipponbare genome was not yet available (until 2005) (International Rice Genome Sequencing Project and Sasaki 2005), and was later improved in 2013 (Kawahara et al. 2013). With the genomes of TN1 (Panibe et al. 2021) and IR8 (Stein et al. 2018) now available, we aim to compare the *sd1* genes of these cultivars and redefine the semidwarf gene based on TN1 and IR8, the two direct descendants of DGWG.

If the greatest strength of TN1 is its high-yielding property due to its semi-dwarf stature from the *sd1* gene, its weakness is its high susceptibility to the blast disease. Rice blast leads to a severe annual loss in rice production worldwide (Wang et al. 2014). However, plants have a natural defense against this and other pathogens, thanks to their resistance genes or R genes. Most R genes are composed of a nucleotide-binding site (NBS) domain and a leucine-rich repeat (LRR) domain (Takken and Joosten 2000). A combination of R genes in a plant may lead to a wide range of immunity response (Fukuoka et al. 2015). Unfortunately, TN1 is susceptible to major rice diseases like blast caused by the fungus *Pyricularia oryzae* (syn. *Magnaporthe oryzae*) (Sabbu et al. 2016) and the bacterial blight disease caused by the bacteria *Xanthomonas oryzae* pv. *oryzae* (Kumar et al. 2012). Predicting the R genes in the genome of TN1 will help understand the resistance profile of TN1, and why it is highly susceptible to blast. For factors that affect plant sensitivity to blast disease, see Chen et al. (2019), Liu et al. (2021), Nugroho et al. (2021) and Zhang et al. (2015).

There are in total 37,526 predicted genes in the TN1 genome (Panibe et al. 2021). Of these thousands of genes, some could be under the influence of positive selection (PS), conferring the cultivar certain advantages that could be related to TN1’s phenotypic characteristics like drought tolerance (Garg and Singh 1971; Garg et al. 2002). Mining the entire genome for genes that makes TN1 unique is no longer highly challenging, thanks to bioinformatics tools that automate the process of looking for positively selected (PS) genes such as PosiGene (Sahm et al. 2017). By using an input of coding sequences from the genomes of GR-related cultivars like IR8 (Stein et al. 2018), MH63 (Zhang et al. 2016) and IR64 (Tanaka et al. 2020) and also other genomes such as maize and wheat, PosiGene may detect the PS genes of TN1.

In brief, this study has three objectives. First, we correct and redefine the *sd1* gene sequence based on the genome assemblies of TN1 and IR8, two direct descendants of DGWG, which is the source cultivar of the semidwarf gene. We then verify the deletions in the *sd1* gene sequence of TN1 and IR8 via Sanger sequencing. We address the questions of which *sd1* sequence fits the previous gene models of the semidwarf gene and “is there really a 383-bp deletion”? To validate the deletion, we compare the *sd1* sequence from the TN1 genome against TN1 reads from the 3,000 Rice Genomes Project (Wang et al. 2018b). Second, we identify R genes in TN1 and investigate why TN1 is highly susceptible to blast, and conduct a haplotype analysis of the blast resistance genes that are apparently missing in TN1. Third, we make full use of the TN1 genome by doing a genome-wide scan to look for PS genes. What are the genes that have undergone positive selection in TN1? What are the functions of these PS genes? Our characterization of the TN1 genome will improve the understanding of the first semi-dwarf cultivar, which initiated the Green Revolution.

## Results

### Defining the regions of the *sd1* gene in TN1 and IR8

Figure 1 shows the portion of the *sd1* gene-to-gene Clustal (Larkin et al. 2007) alignment showing the first 427 bp of TN1, 518 bp of IR8, 890 bp of Nipponbare, and 426 bp sequence by Monna et al. (2002). The 383 bp deletion discovered by Monna et al. (2002) became 382 bp in TN1 and IR8 because of the adenine at position 285 and position 376, respectively. The alignment continues in Appendix, Figure S1. All positions refer to chromosome 1. OsTN1g004133 position 1 is 40,361,934. OsIR8_01G0407900 position 1 is 39,824,196. Os01g0883800 position 1 is 38,382,466. The deleted region in Nipponbare *sd1* lies between 38,382,762 and 38,383,144 of the genome.

**Figure 1.**
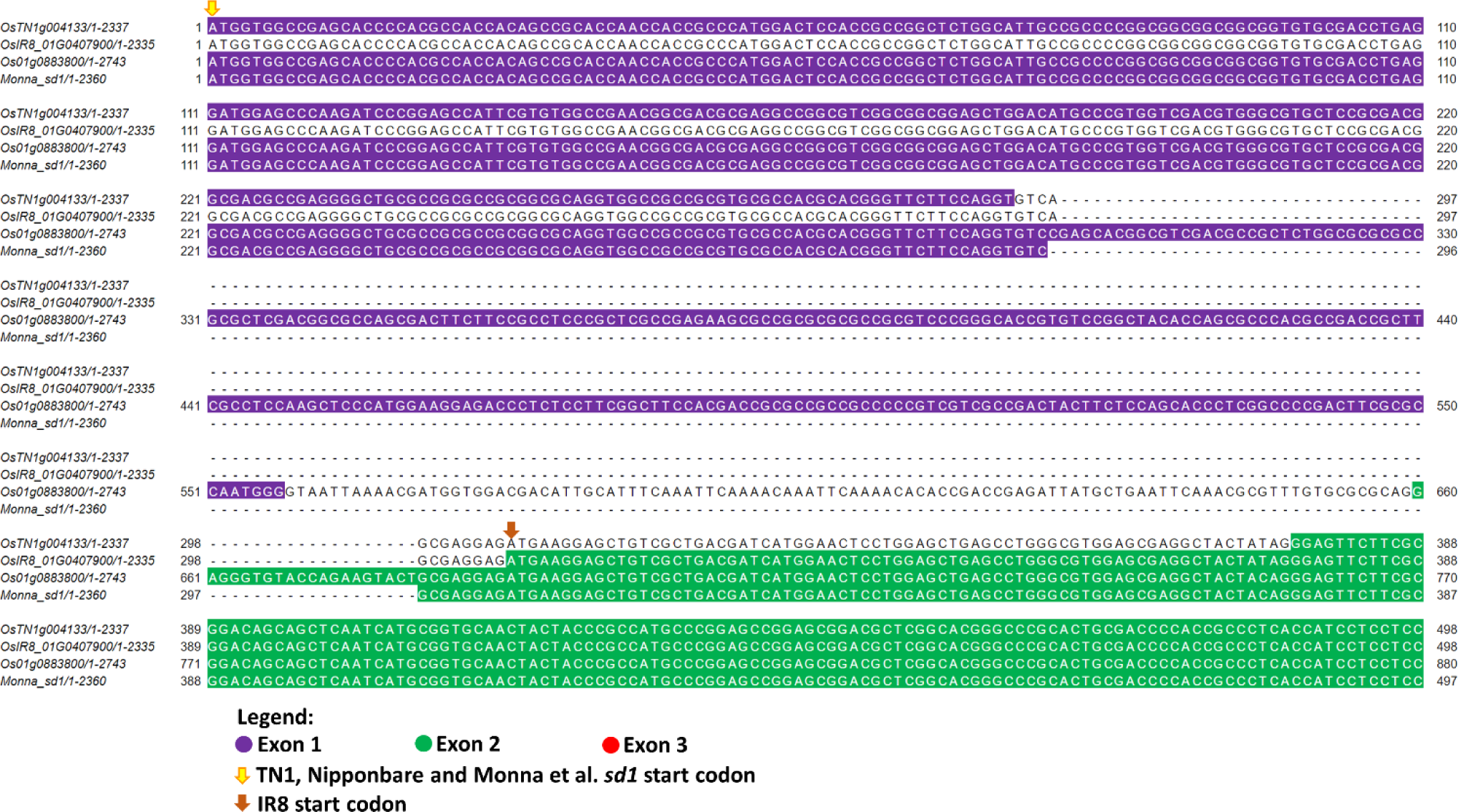
Portion of the Clustal alignment of the *sd1* exon sequences in TN1, IR8 and Nipponbare. Position 1 is the first nucleotide of exon 1 of the gene. For TN1, IR8 and Nipponbare, position 1 is located at 40,361,934; 39,824,196; and 38,382,466, of chromosome 1, respectively. OsTN1g004133, OsIR8_01G0407900 and Os01g0883800 refer to the *sd1* gene IDs of TN1, IR8 and Nipponbare, respectively. Beside the gene IDs are the positions of the first base and last base of the sequence, with the latter representing its total length. The total sequence length as well as the start and end positions of each exon are depicted in Figure 2. Deleted nucleotides are represented by dashes. The location of the first base of the start codon is indicated by an arrow pointing downwards. Exons are colored whether they are the first, second or third exon. Nucleotides in introns are uncolored, except for the sequence before the start codon of IR8, which was assigned as a 5’ untranslated region in its annotation.

To better understand the *sd1* gene of TN1 and IR8, we show their gene structure models derived from protein sequence alignment and gff annotation (Gramene 2020; Panibe et al. 2021), and compared it to Nipponbare (Figure 2). TN1 has 3 exons and 2 introns. It has an exon gap that spans half of Nipponbare’s exon 1 up to one-third of exon 2 of the *japonica* cultivar (see Figure 2a). The exon gap in TN1 *sd1* does not represent the 382 bp deletion but rather the lost coding sequence as defined by its gff annotation. For IR8, its exon 1 seems to become lost due to an untranslated region (UTR) as indicated in its gff annotation.

**Figure 2.**
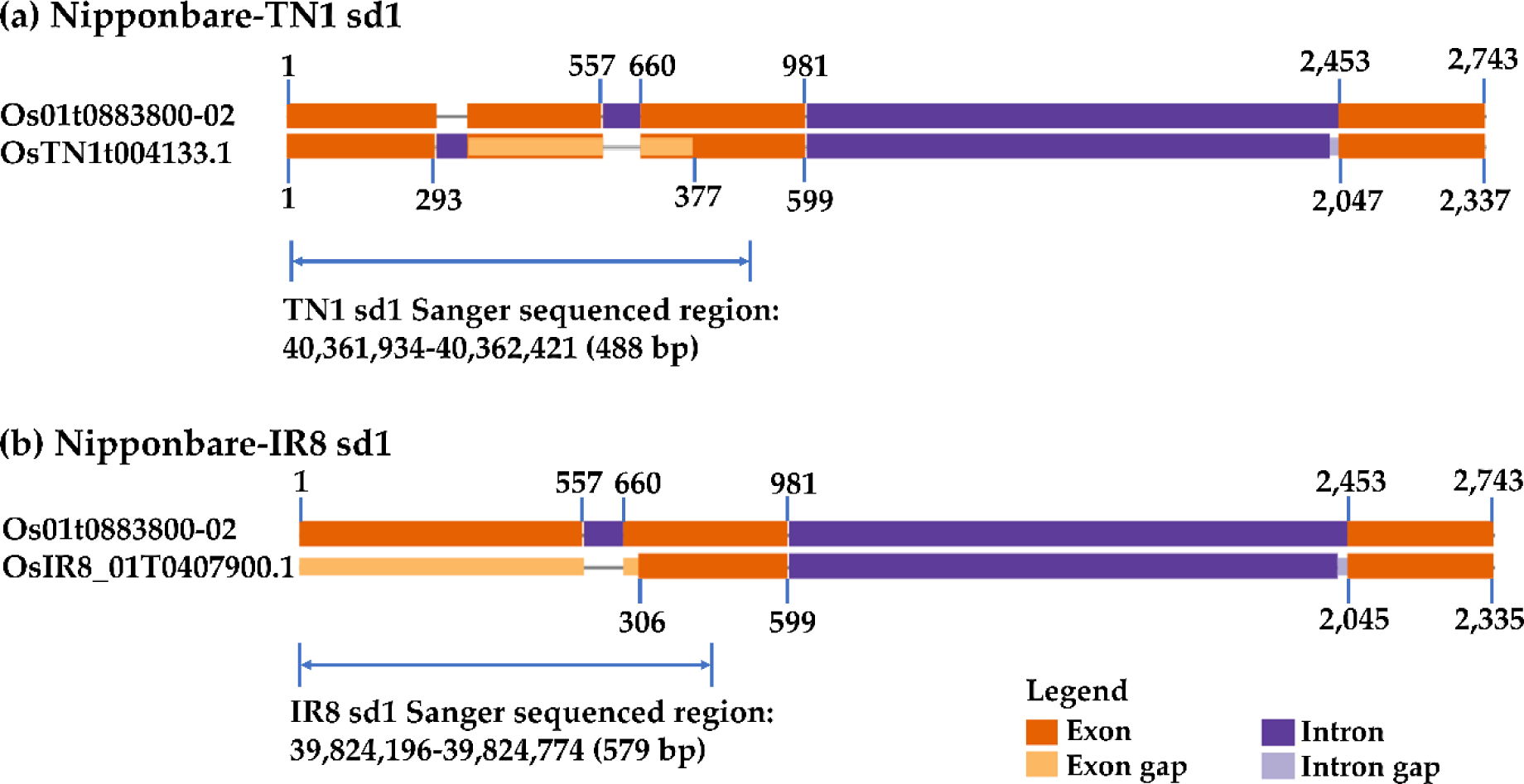
The *sd1* exon to exon gene diagrams of TN1, IR8 and Nipponbare. TN1 (2,337 bp), IR8 (2,335 bp), and Nipponbare (2,743 bp). (a) Nipponbare vs TN1 and (b) Nipponbare vs IR8. Os01t0883800-02 is from Nipponbare, OsTN1t004133.1 is from TN1, and OsIR8_01T0407900.1 is from IR8. These names are based on the transcript ids of their *sd1* gene. Numbers indicate the exon boundaries of the genes. For TN1 position 1 is at 40,361,934 bp, while for IR8 position 1 is at 39,824,196 bp. In Nipponbare, position 1 is at 38,382,466. The gene diagrams were created by GenePainter (Hammesfahr et al. 2013) by using the protein alignments of *sd1* and the information from their gff annotation (Nagano et al. 2005; Panibe et al. 2021). The range specified by the light blue arrow represents the sequences of *sd1* in TN1 and IR8 that were validated by our Sanger sequencing. The 382 bp deletion in TN1 can be derived by computing the difference between 981 and 599, the latter of which represents the gene length of TN1 *sd1* before its 2nd intron.

We further confirmed the *sd1* gene sequence of TN by mapping TN1 short reads used in the 3000 Rice Genomes Project (3K RGP) (Wang et al. 2018b). There are actually two sets of TN1 reads in the 3000 Rice Genomes Project and they have the assay IDs, CX270 and CX162. The former has the name TAICHUNGNATIVE1, while the latter is designated as TN1. To determine which one better represents the sequencing reads from the 3000 Rice Genomes Project, we mapped the reads to the TN1 genome. CX162 has a 99.92% and 90.92%, for the overall mapping rate and properly paired mapped reads. respectively. In contrast, CX270 has a mapping rate of 99.40% and 81.91%. Based on the mapping of reads, CX162 better represents the TN1 genome in the 3K RGP.

We also checked the SNP-Seek database (Mansueto et al. 2017), if there are SNP loci inside the region corresponding to the *sd1* deleted sequence in semi-dwarf cultivars. Of the two, TN1 (CX162) has missing SNP positions to deletion in *japonica* (Figure 4a), whereas TAICHUNGNATIVE1 (CX270) has alleles on the same set of coordinates. (Figure S3). We further inspected the mapping of the reads by viewing the *sd1* region in Integrative Genomics Viewer (IGV) (Robinson et al. 2011), and they are shown in Figure 4b (CX162) and Figure S4 (CX270). The nucleotide at chromosome 1 position 40,362,230 was supported by the TN1 reads of CX162 (Figure 4c) and CX270 (Figure S5). The former’s reads better covered the position compared to the latter. In CX162, it is mapped by six reads, while in CX270 it is by only one read.

**Figure 3.**
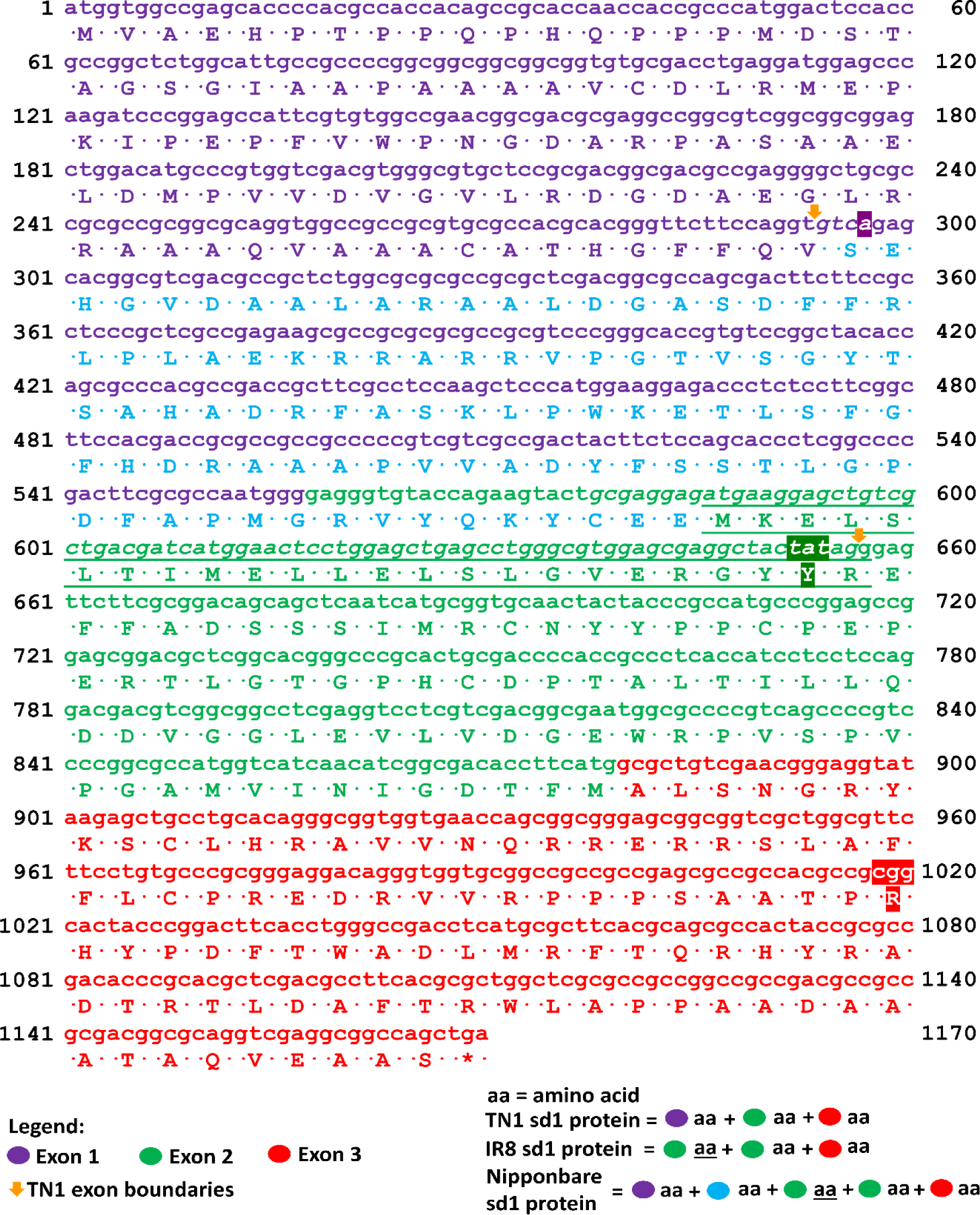
Predicted coding and amino acid sequences of Nipponbare, TN1 and IR8 *sd1* gene sequences. Position 1 is 40,361,934; 39,824,196; and 38,382,466 in chromosome 1 of TN1, IR8 and Nipponbare, respectively. The IR8 *sd1* sequence illustrated here is based on its gff annotation dated May 10, 2020 and not yet corrected. TN1 and IR8 CDS and protein sequences were aligned to the Nipponbare CDS via Clustal to see the similarity of the sequences per position. The same Nipponbare CDS was translated and the mapping of its amino acids per codon became the basis of this diagram, such that all nucleotides and amino acid translations were the same for the three cultivars unless indicated. The nucleotides with amino acid translation that are colored light blue and not italicized are the deleted CDS region in TN1 *sd1*, which is the exon gap in Figure 2a. The italicized nucleotides (including the violet shaded adenine) is intron 1 of TN1 *sd1*. The colored codon tat (TN1 and IR8) is the synonymous codon of tac of Nipponbare, while codon cgg (TN1 and IR8) is a mutation for codon cag (Nipponbare), which changes amino acid Q (Nipponbare) to R (TN1 and IR8). The violet shaded adenine in codon 1 is the extra nucleotide that caused the 1 bp difference of the genomic deletion of TN1 and IR8 against the *sd1* sequence of Monna et al. (2002).

**Figure 4.**
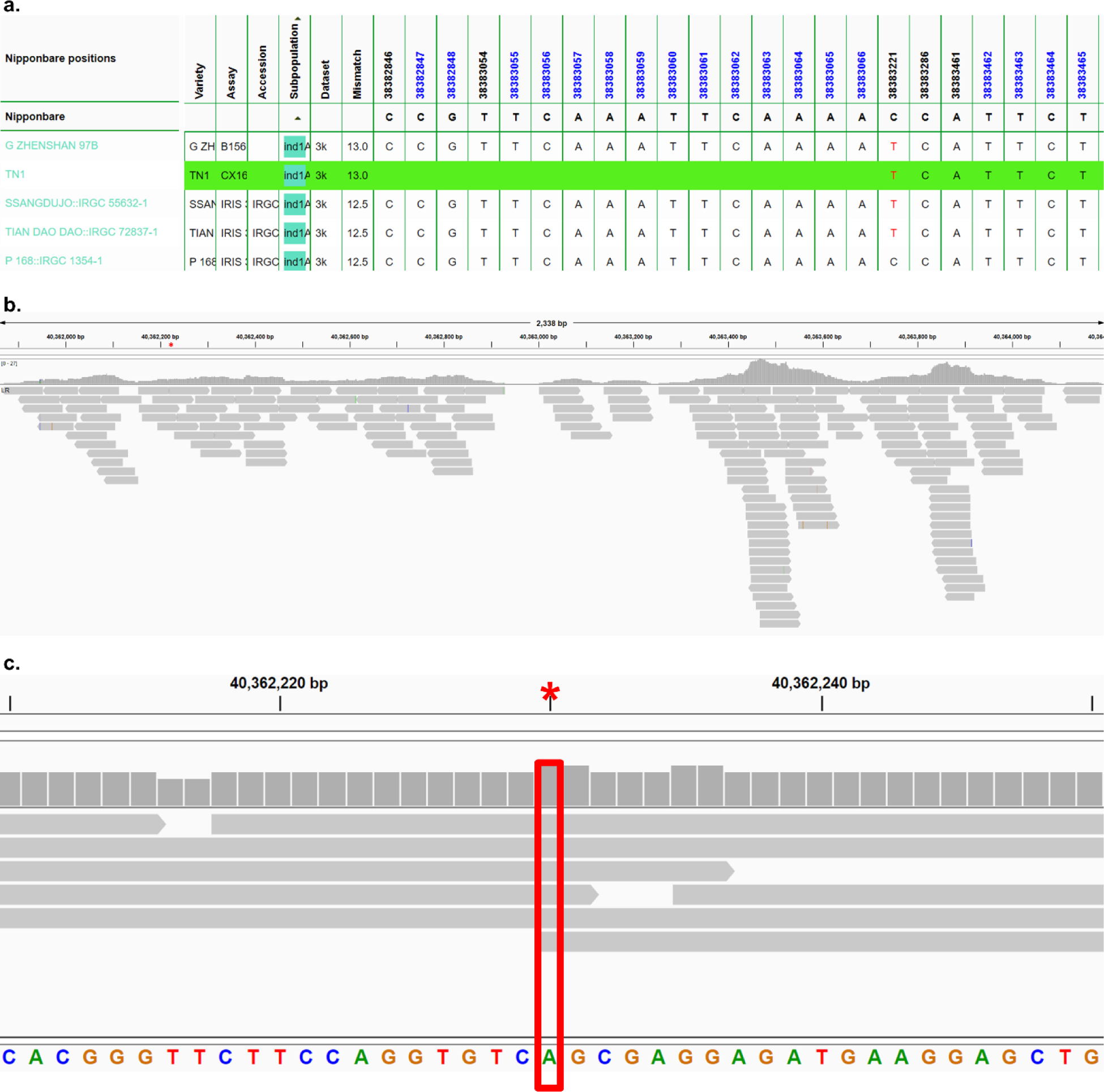
*sd1* gene SNP-Seek result for TN1 and mapping of the CX162 reads. a) Screenshot of the Genotype search result of the SNP-Seek database for the *sd1* gene. The blank green space for the TN1 cultivar assay CX162 signifies the absence of genomic DNA in TN1 with respect to the Nipponbare reference genome. According to the alignment in Figure 1, the deleted region in Nipponbare *sd1* lies in between 38,382,762-38,383,144 of chromosome 1 and is consistent with the missing TN1 SNPs in SNP-Seek with respect to the Nipponbare reference genome starting at position 38,382,846 up to position 38,383,066. b) Mapping coverage of the TN1 (assay CX162) 3K RGP reads onto the *sd1* gene region of the TN1 genome. The bam file was filtered to only get the properly paired reads with the proper distance. The gray color means that the read sequence is identical to the TN1 genome. The red asterisk is the approximate location of the start of the *sd1* deletion with respect to the Nipponbare genome. c) Closeup of the asterisk area in Figure 4b. It has the coordinate 40,362,230 in TN1 chromosome 1. It corresponds to the extra nucleotide that made the *sd1* deletion in TN1 and IR8 382-bp, instead of 383-bp. It is covered by 6 properly mapped paired-end reads with the correct distance. Gray color means the read sequence are homozygous to the TN1 genome.

### Predicted R genes in TN1

We annotated 383 NLR (nucleotide-binding domain leucine-rich repeat), 34 NB-ARC and 6 LRR (leucine-rich repeat) in the TN1 genome (Dataset S1). For this purpose, we used the Tetep as a reference because Tetep is known to be highly resistant to blast disease and its genome and R genes have been well characterized (Wang et al. 2019b); indeed, it has been commonly used to breed for new blast resistant cultivars (Singh et al. 2012; Zarbafi and Ham, 2019; Ramalingam et al. 2020). The numbers of orthologues found between Tetep (Wang et al. 2019b) and TN1, MH63, R498 and Nipponbare did not show significant differences (Appendix, Table S1). Non-orthologous Tetep NLRs (R genes) were then blasted against the TN1 proteome using their NR-ARC domain protein sequences (Dataset S2) and those hits with alignment identity <50% were deemed missing in the TN1 assembly. One of the unfound R genes in TN1 is *Pi54* (*Pik-h*), which is the gene chr11.fgenesh2107 in the assembled Tetep genome (Wang et al. 2019b). *Pi54*, originally cloned from Tetep, is known to confer broad-spectrum resistance to blast (Gupta et al. 2011; Rai et al. 2011; Thakur et al. 2015). Moreover, ∼28 of the 90 NLR genes that were found to be resistant to one or more blast fungal strains (Wang et al. 2019b) were found to be missing or mutated in the TN1 genome (Dataset S2).

By using the method of Mahesh et al. (2016), the set of 22 cloned blast R genes were searched in the TN1 genome. The results are given in Table 1 and those marked with an asterisk were the results different from Mahesh et al. (2016). These genes are confirmed to be present by Blastp in the Tetep genome with the same criteria used by Mahesh et al. (2016), i.e., e-value < 10e-10, identity ≧ 70% and query coverage ≧ 70% (Dataset S3). The same set of R genes were also searched in the TN1 genome. Some of the R genes are present in TN1 but are mutated (Table 1), preventing the translation of the gene into the right protein. In the case of Tetep, both Wang et al. (2019b) and Mahesh et al. (2016) found the *Pi-ta* and *Pi54* R genes in the blast resistant cultivar (see the Tetep column in Table 1).

**Table 1.**
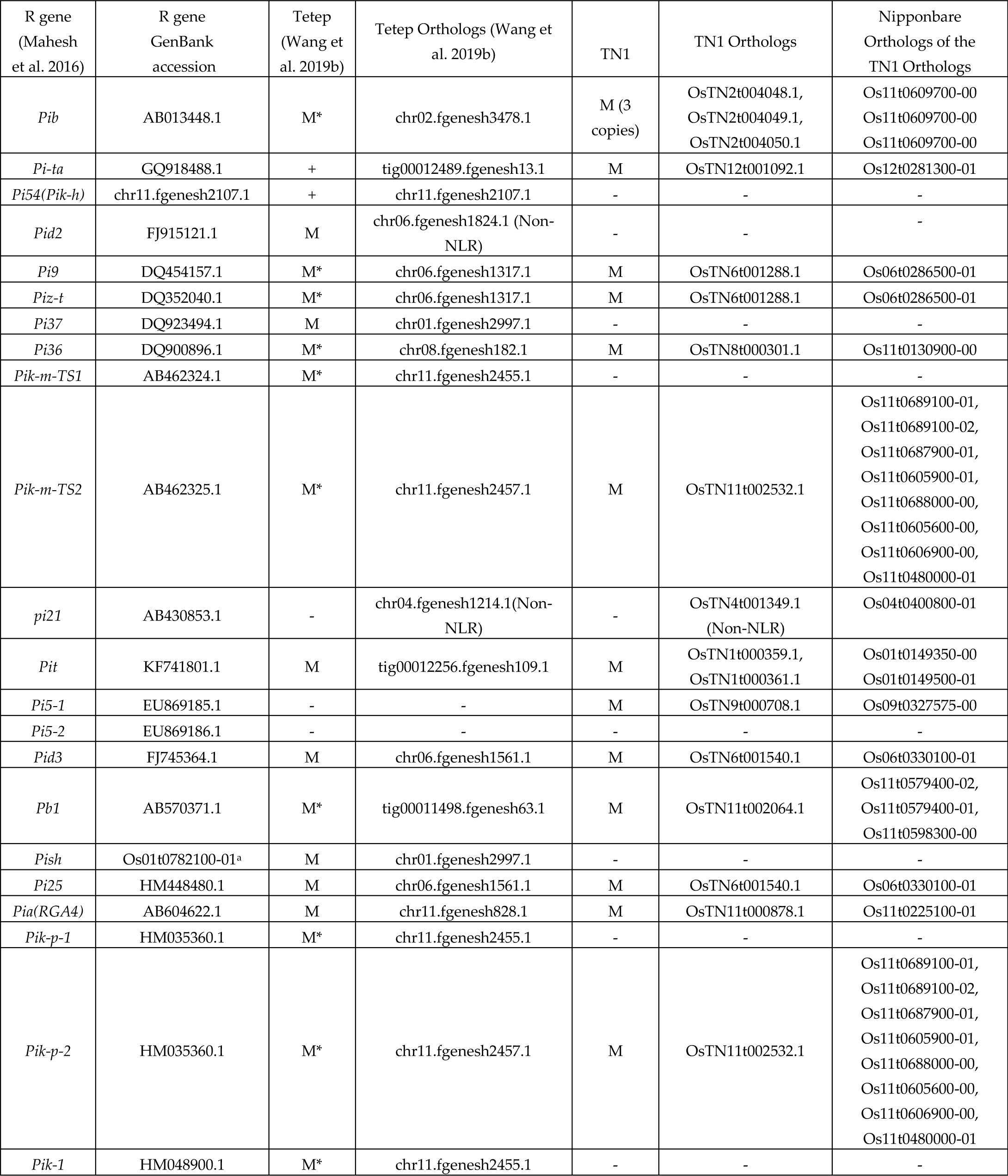
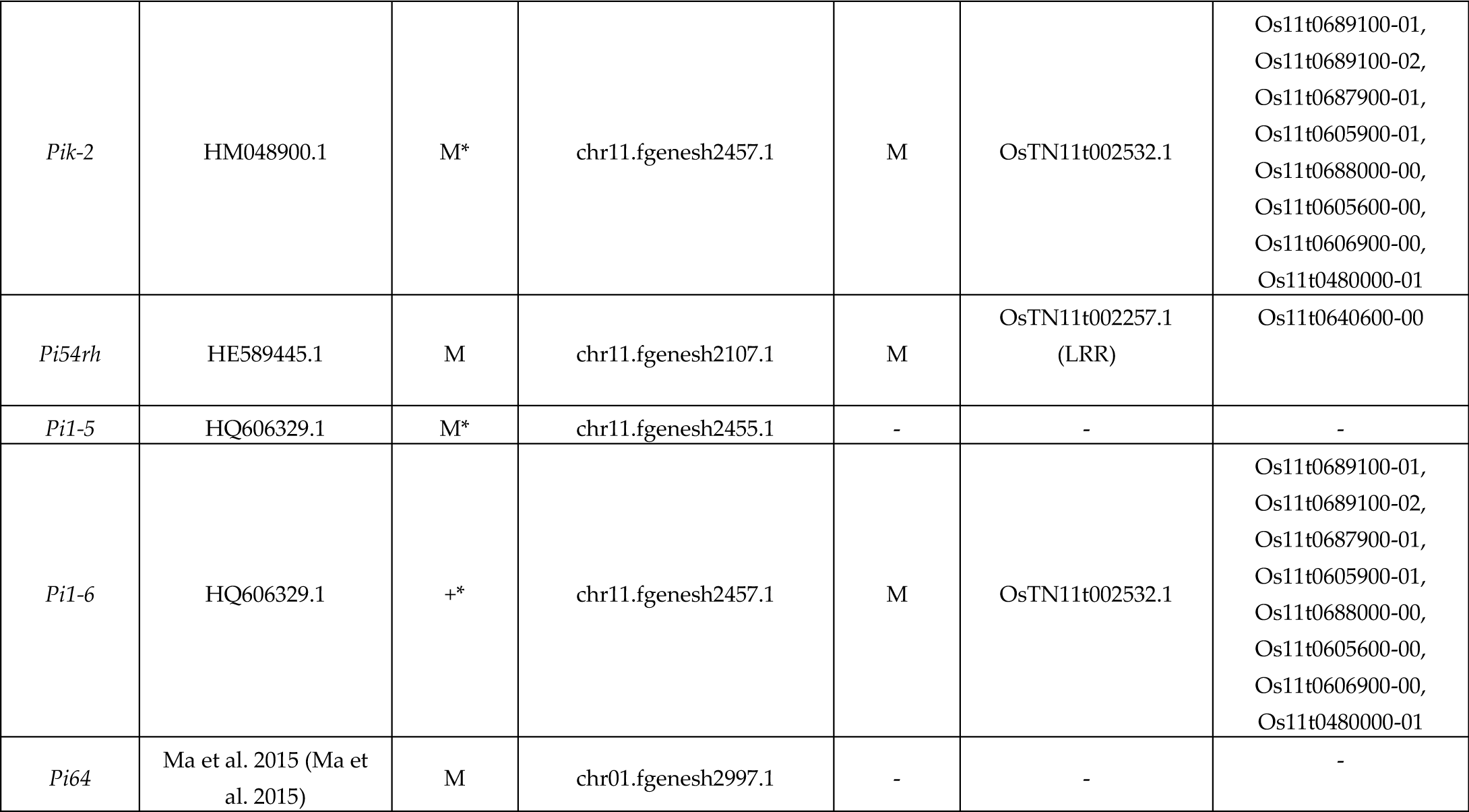
Distribution of cloned blast resistance genes in sequenced rice varieties. A + means present and a - means absent while M means mutated but with protein structure retained. * means a result different from Mahesh et al. (2016). These genes are confirmed to be present by blastp in the Tetep (Wang et al. 2019b) genome with e-value < 10e-10 and identity > = 70% (see Dataset S3 for details). chr11.fgenesh2107.1 is a gene name from the Tetep genome annotation.

### Haplotype analysis of the *Pi-ta* and *Pi54* genes

To filter the missense variants, we compared each allele in the haplotypes of *Pi54* and only obtained the SNPs that are heterozygous (see Methods). Only three SNP positions were left and they are located on chromosome 11: 25,263,636; 25,264,119; and 25,264,164 (see Table 2). We checked their allele frequencies and found that the missense variants have a minimum allele frequency (MAF) of 36% to 39% (Table 2), suggesting that these missense variants are maintained across the rice populations, even though each causes a change in amino acid. Using the major/minor allele section in Table 2, we compared the three alleles of TN1 to the major and minor alleles. At SNP position 25,263,636, TN1 has two possibilities: either allele C or T (Table 2). If it is a T, it will be a minor allele across the 3,024 rice cultivars. The missense variant causes a Glu144Lys mutation (Dataset S5), changing an acidic amino acid into a basic one. The change in charge of an amino acid could disrupt ionic interactions in the structure of the protein, which could affect its function, supporting our observation from Table 1 that the *Pi54* gene in TN1 is missing when compared to the blast resistant Tetep cultivar.

**Table 2.** *Pi54* alleles of blast susceptible and resistant cultivars at the same SNP position. Haplotype numbers are based on the kgroup numbers in Dataset S4. SNP information and allele frequency were obtained from SNP-Seek (Mansueto et al. 2017). The reference genome is the Nipponbare cultivar, which was included in the study of Thakur et. al (Thakur et al. 2015) and also part of the 3K RGP. Alleles of Tetep were obtained through show-snps of the MUMmer version 4 package (Marçais et al. 2018), after nucmer alignment of Tetep chromosome 11 (which has the *Pi54* gene) against Nipponbare chromosome 11.

| Cultivar | Resistant or Susceptible | # mismatches | Alleles at chromosome 11 |  |  |
| --- | --- | --- | --- | --- | --- |
|  |  |  | SNP position |  |  |
|  |  |  | 25,263,636 | 25,264,119 | 25,264,164 |
| Nipponbare | Susceptible<br>(Thakur et al.<br>2015) | 0 | C | G | T |
| CO-39 | Susceptible<br>(Thakur et al.<br>2015) | 12 | C | A | C |
| Parijat | Susceptible<br>(Thakur et al.<br>2015) | 0.5 | C/T | G | T |
| Pusa basmati 1 | Susceptible<br>(Thakur et al.<br>2015) | 24 | C | A | C |
| Budda | Resistant<br>(Thakur et al.<br>2015) | 30 | C | A | C |
| IRAT-144 | Resistant<br>(Thakur et al.<br>2015) | 4.5 | C/T | G/A | T/C |
| Salumpikit | Resistant<br>(Thakur et al.<br>2015) | 2 | T | G | T |
| Tetep | Resistant<br>(Wang et al.<br>2019b) | - | C | A | C |
| TN1 | Susceptible | 1.5 | C/T | G | T |
| Haplotype |  |  |  |  |  |
| Haplotype 1 | - |  | T | G | T |
| Haplotype 2 | - |  | C | A | C |
| Major/Minor allele<br>and MAF |  |  |  |  |  |
| Major allele | - |  | C | G | T |
| Minor allele | - |  | T | A | C |
| Minor allele<br>frequency (MAF) | - |  | 38.80% | 35.86% | 35.84% |

We also investigated the *Pi-ta* gene in SNP-Seek and it returned four haplotypes (Dataset S4). This is the same number of haplotypes that Jia et. al (Jia et al. 2003) found. In that study, three of the haplotypes were related to susceptibility to blast and have five nucleotide positions that caused a non-synonymous mutation in *Pi-ta*. We checked the annotation of the SNPs in the 3K RGP and found that the I6S mutation (Dataset S5) was due to the replacement of the G nucleotide by a T at position 10,611,754 (Table 3). TN1 has the A allele at this position, which is the minor allele across the 3,024 cultivars in SNP-Seek. Consequently, TN1 is predicted to have the I6S mutation in its *Pi-ta* protein. From Table 3, the resistant cultivars Katy and Drew have the alleles T,G and C at positions 10,611,244; 10,611,297; 10,611,327, and an A at position 10,611,754.

**Table 3.**
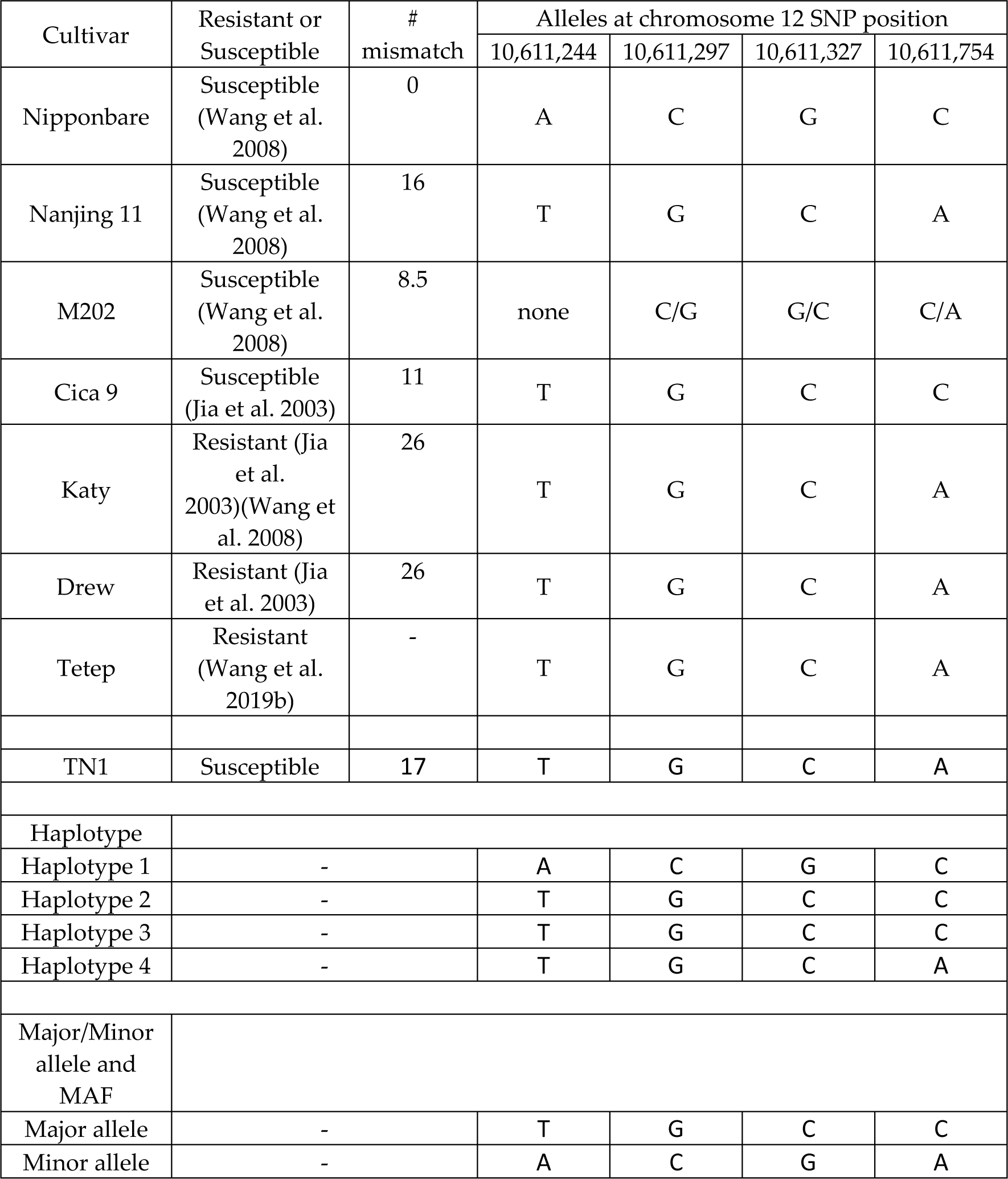
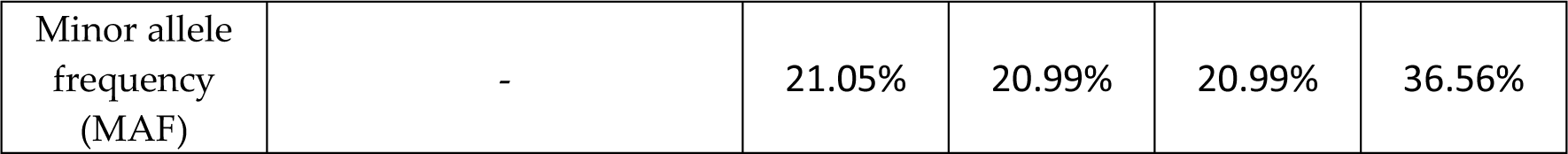
*Pi-ta* alleles of blast susceptible and resistant cultivars at the same SNP position. Haplotype numbers are based on the kgroup numbers in Dataset S4. SNP information and allele frequency were obtained from SNP-Seek (Mansueto et al. 2017). The reference genome is the Nipponbare cultivar, which is susceptible according to Jia et. al (Thakur et al. 2015) and Wang et. al (2008). It is also part of the 3K RGP. Alleles of Tetep were obtained through show-snps of the MUMmer version 4 package (Marçais et al. 2018), after nucmer alignment of the Tetep contig tig00012489 (which has the *Pi-ta* gene) against Nipponbare chromosome 12.

However, the susceptible Nanjing 11 cultivar as well as TN1 has the pattern of alleles at the mentioned SNP positions similar to Katy and Drew. We did not see very clear difference between the haplotypes of the susceptible ones and the resistant ones for *Pi-ta* (Table 3) but for *Pi54* the differences look like a bit clearer (Table 2). *Pi54* is considered non-functional as the allele in TN1 (OsTN11t002257.1) lost the first 598 amino acids when compared to Tetep (Figure S6), resulted in complete loss of the NB-ARC domain. Of the 9 absent cloned NLRs (11 alleles shown in Table 1, which belong to 9 genes) in TN1, only 3 are from *indica* donors (other 6 might represent *japonica*/*indica* differences), including *Pi54*, *Pid2* and *Pi1-5*. *Pid2* is present in both susceptible and resistant cultivars, while Pid1-5 is absent in them all. Only *Pi54* shows presence/absence polymorphism in resistant (Tetep and Tadukan) and susceptible (Co-39 and HR-12) cultivars (Mahesh et al. 2016). We further investigated the *Pi-ta* gene of TN1 by aligning it against its counterpart in Yashiro-mochi (a resistant cultivar). The protein sequence alignment of *Pi-ta* in TN1 is largely the same compared to the latter (Figure S7), suggesting that the function of the gene in TN1 is not largely altered.

### Eleven genes in TN1 underwent positive selection

The aim of the genome-wide search for TN1 genes that underwent positive selection (PS) is to identify genes that might explain TN1’s phenotypic characteristics like high yield (Yoshida 1981), photoperiod insensitivity (Vergara and Chang 1985) and drought-tolerance (Garg and Singh 1971). The GO terms assigned to these PS genes would give insights into the biological processes involved as well as the enzymes that confer the function. We identified 11 TN1 genes that were likely subject to PS in TN1 in the past (Table 4).

**Table 4.** The 11 TN1 genes that underwent positive selection. . The criterion for positive selection was FDR <0.05. The PS sites are indicated in the CDS alignments created by PosiGene in Appen-dix, Figure S10.

| Gene | Description | FDR | # of PS sites |
| --- | --- | --- | --- |
| OsTN5g000040 | hypothetical protein | 6.23E-11 | 5 |
| OsTN5g002486 | - | 1.93E-05 | 3 |
| OsTN2g002903 | PLATZ transcription factor family protein | 3.73E-05 | 1 |
| OsTN5g001087 | GA 3β-hydroxylase | 4.12E-05 | 2 |
| OsTN1g003572 | armadillo/beta-catenin repeat protein-like | 4.12E-05 | 2 |
| OsTN1g003413 | transmembrane protein 56 isoform X1 | 5.07E-05 | 2 |
| OsTN8g001161 | probable TPP C | 5.07E-04 | 4 |
| OsTN12g002058 | L-type lectin-domain containing receptor<br>kinase IX.1-like | 1.55E-03 | 2 |
| OsTN12g001576 | - | 1.88E-03 | 1 |
| OsTN1g000744 | KARI, chloroplastic | 3.50E-02 | 1 |
| OsTN4g000152 | probable LRR receptor-like<br>serine/threonine-protein kinase<br>At3g47570 | 4.90E-02 | 2 |

Using the Blast2GO annotation of the TN1 assembly (Panibe et al. 2021), a total of 35 GO terms (Appendix, Table S3) were assigned to six of the 11 PS genes (Table 4); see their representative GO terms in Figure 5. For the Molecular Function (Figure 5b), a correlation is observed between the protein names of the six genes and their GOs.

**Figure 5.**
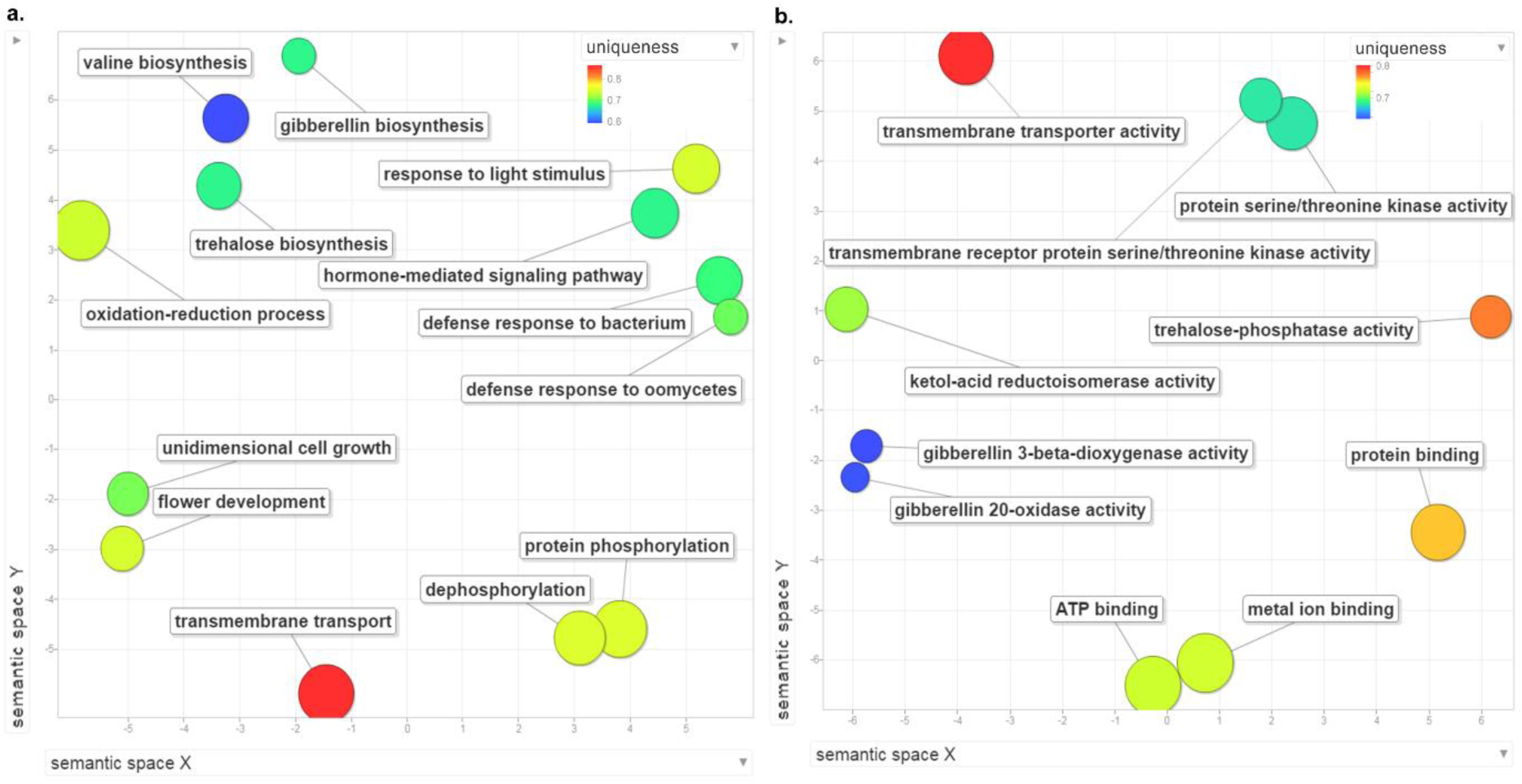
REVIGO (Supek et al. 2011) visualization of GOs of genes under positive selection in TN1. a) Biological Process and b) Molecular Function. These are the scatterplot of REVIGO showing the representative GO terms of the PS genes, where the colors of the circles represent the uniqueness value, computed from comparing each GO term to each other. The bigger the size of the sphere, the more the general the term is.

## Discussion

### *sd1* has a 382-bp deletion in the semidwarf TN1

To redefine the *sd1* gene, we first compared the *sd1* genes of TN1, IR8, Nipponbare and the sequence by Monna et al. (2002) (Figure 1). The alignment of TN1 and IR8 shows a 382 bp deletion, in contrast to Monna et al.’s (2002) 383 bp deletion (Figure 1). The same observation was found in the *sd1* gene of the parent DGWG (Nagano et al. 2005) cultivar, as well as two of its indirect descendants, MH63 (Wu et al. 2017; Jia et al. 2020) and IR36 (Jia et al. 2020). The presence of an adenine nucleotide at position 285 of TN1 and at position 376 of IR8 was the reason for the difference (Figure 1). Further inspection of the alignment (Appendix, Figure S1) shows that there are point mutations and deletions between the TN1/IR8 and Monna et al.’s (2002) sequence. With TN1 and IR8 as the direct descendants of DGWG, the *sd1* genes from these two cultivars should better represent the gene than Monna et al.’s (2002) sequence, which was derived from DGWG-type *sd1* mutants (Habataki, Milyang 23, and IR24) (Monna et al. 2002).

Thus, TN1’s *sd1* exon 1 encodes up to 98 amino acids only because of its exon-intron boundary (Figure 3). The fact that the “missing” Sanger validated sequence of IR8 is identical to TN1 suggests that the two cultivars have the same protein sequence. The confusion in the length of the coding sequence deletion (280 bp in this study, 280 bp in (Spielmeyer et al. 2002), and 278 bp in (Monna et al. 2002)) is clarified if the *sd1* annotation of TN1 is used, and the Green Revolution *sd1* sequence refers to TN1 only. To correct the IR8 *sd1* sequence, we suggest to use the CDS and amino acids of TN1 as shown in Figure 3. To sum up, the coding and protein sequence of the TN1 and IR8 cultivars are identical, if the annotation of TN1 *sd1* is used.

We validate the deletion in the *sd1* gene sequences of TN1 and IR8 by Sanger sequencing. To get a fair comparison of the differentiated region, we sequence the region from the first nucleotide of exon 1 of TN1 and IR8 *sd1* up to one-half the length of exon 1 of the semidwarf cultivars before the exon-intron boundary corresponding to position 981 of the Nipponbare gene structure; see blue arrows in Figure 2. For TN1, the coordinates are chr 1:40,361,934-40,362,421 (Figure 2a). For IR8, the region validated includes the 5’ untranslated region (UTR) (chr 1: 39,824,196-39,824,774) because that is part of IR8’s *sd1* exon 1 as indicated in its gff annotation (Gramene 2020). In Figure 2b, the 5’ UTR has become part of the exon gap. To validate that the differentiated regions really exist, we align via Clustal the Sanger sequences to the nucleotides extracted from the genome assemblies of TN1 and IR8. The resulting alignment shows that the *sd1* sequence of TN1 is 100% identical to and 100% covered by the Sanger sequences (Appendix, Figure S2). Likewise, the IR8 *sd1* sequence from its genome matches its Sanger sequence. When the two Sanger sequences are compared, the TN1 has a perfect overlap with its IR8 counterpart, covering its entire 488 bp length. Because both TN1 and IR8 derived their semidwarf gene from their parent DGWG, the two cultivars should have the same form of the *sd1* gene. This suggests that the untranslated region in the IR8 *sd1* defined by its annotation is not really a UTR region, but an exon-intron-exon structure similar to TN1 (Figure 2a). In lieu of this, we propose that the gene model of IR8 should follow that of TN1 and that the current annotation of the IR8 *sd1* gene is in error.

We also compare the coding sequences of TN1, IR8 and Nipponbare. Spielmeyer et al. (2002) reported that the first 99 amino acids from the CDS of semidwarf Doongara, a descendant of DGWG, is similar to that of the Nipponbare, and that there is a 280 bp deletion in the coding sequence. Meanwhile, Moona et al. (2002) reported a 278 bp deletion in the expressed sequence of DGWG-type cultivars. The alignment in Figure 3 indicates that there is 280 bp deletion in the coding sequence of TN1. We obtain this number by computing the difference between the length of the deletion (363 bp) in Figure 3 and the length of first intron of TN1 *sd1* (83 bp). There is also a frameshift mutation in the CDS of TN1 *sd1* but this occurs at the junction of position 293 and position 294 (Figure 3). However, the codon does not change because of the same guanine nucleotide at the start of exon 2, leading to the same valine amino acid.

### The *Pi54* resistance gene in TN1 is missing

TN1 is known to be highly susceptible to the blast fungus and the cultivar was used as a standard in searching for resistance genes (Sabbu et al. 2016). Using the SES (Standard Evaluation System) for Rice (International Rice Research Institute 2013), which designates a score of 0 to 9 with increments of 1 for the varying severity of the blast disease caused by Pyricularia oryzae. A score of 0 (no spots) to 1 (tiny dots) is considered highly resistant and score of 8 to 9 means highly susceptible (International Rice Research Institute 2013). The score is based on the size of the area damaged by the pathogen on the leaves. TN1 was given a score of 9, wherein 75% of the leaves succumb to P. oryzae, while Tetep was assigned a score of 1 against the blast fungus (Sabbu et al. 2016). Tetep harbors the R genes *Pi-ta* (Mahesh et al. 2016; Wang et al. 2019b), *Pi54*(*Pik-h*) (Sharma et al. 2005), and *Pitp(t)* (Barman et al. 2004). Thus, Tetep is a good reference in searching for blast R genes in TN1. From the list of predicted R genes in TN1, we looked for the orthologues of TN1 R genes in Tetep and catalogued any mutations between the orthologues. We narrowed down the list of blast R genes to check by using the set of resistance genes studied by Mahesh et al. (2016).

Of the two genes, we suspect the direct absence of *Pi54* in TN1 (Table 1) to partly cause its blast susceptibility. The logic is simple: (1) we analyzed nearly all best functionally studied NLR genes in rice (i.e., the 22 genes), and only *Pi54* shows presence/absence polymorphism between *indica* resistant (e.g., Tetep and Tadukan) and susceptible (e.g., HR-12 and Co-39) cultivars and is absent in TN1 (Table 1); (2) *Pi54* confers broad spectrum resistance to blast disease, and is being used in some enhanced blast resistant breeding programs (Thakur et al. 2015). Haplotype analysis of 92 cultivars for the *Pi54* gene revealed one haplotype out of 50, called H_3, that is composed of blast resistant *indica* cultivars (Thakur et al. 2015). We expanded the haplotype analysis for *Pi54* by checking the SNP-Seek database (Mansueto et al. 2017), which contains data from pre-computed analysis of 3,024 rice cultivars aka the 3,000 Rice Genomes Project (Wang et al. 2018b). However, instead of getting 50 haplotypes, the alleles of the 3K RGP were grouped to only two haplotypes (Dataset S4). Seventeen SNPS were missense variants (Dataset S5).

### Functions of the genes subjected to positive selection in TN1

For GA 3β-hydroxylase, it is gibberellin 3-beta-dioxygenase activity (GO:0016707). Probable TPP (trehalose-phosphate phosphatase) C has the function of trehalose-phosphatase activity (GO:0004805), while KARI, chloroplastic for ketol-acid reductoisomerase activity (GO:0004455), is involved in biosynthesis of branch chain amino acids valine (GO:0009099) and isoleucine (GO:0009097). For the transmembrane transporter activity (GO:0022857), it refers to the transmembrane protein 56 isoform X1 gene.

TPP (EC:3.1.3.12) and trehalose-6-phosphate synthase (TPS) (EC:2.4.1.15) are important enzymes in trehalose biosynthesis. TPP acts on the product of TPS, which is trehalose-6-phosphate (T6P), dephosphorylating it to produce the end-product trehalose, a disaccharide composed of two glucose molecules linked by an α(1→1) glycosidic bond.

Trehalose is a non-reducing sugar (Stick and Williams 2009), stable enough to become a natural anti-desiccant (Luyckx and Baudouin 2011). This property of trehalose was studied in a fusion gene of TPS and TPP in transgenic rice that led to an increase in trehalose, inducing the plants to become resistant to drought, sodicity and low temperatures (Garg et al. 2002). T6P has been associated with increased yield. In wheat, an increase in T6P led to an increase in yield through the inhibition of sucrose nonfermenting 1 (SNF1)-related protein kinase 1 (SnRK1), while in maize a decrease in T6P led to increased activity of SnRK1, leading to more sucrose transport and an increase in yield (Paul et al. 2018). TN1 is reported to be a drought-resistant cultivar (Garg and Singh 1971) as well as a high-yielding variety. This suggests that the OsTN8g001161 PS gene encoding for TPP could have played a role in this drought resistant, high-yield characteristic of TN1, either by an increase/decrease in T6P or through an enhanced production of trehalose.

The two GO terms protein serine/threonine kinase activity (GO:0004674) and transmembrane receptor protein serine/threonine kinase activity (GO:0004675) are synonymous to each other, and they refer to two different proteins, probable LRR receptor-like serine/threonine-protein kinase At3g47570 and L-type lectin-domain containing receptor kinase IX.1-like. The former is a type of leucine-rich repeat receptor like kinase (LRR-RLK), while the latter is commonly called LecRK or a lectin receptor kinase. The 309 LRR-RLKs in Nipponbare (Sun and Wang 2011) have a role in abiotic stress response (Dievart et al. 2016), while LecRKs are associated with plant immunity (Wang and Bouwmeester 2017). The BP GO terms (Appendix, Table S3) of defense response to oomycetes (GO:0002229) and defense response to bacterium (GO:0042742) support the notion of stress response to pathogens through the LecRK PS gene. However, the LecRK in TN1 could have other functions. In Nipponbare, OsLecRK is not only involved in immune response but also in seed germination (Cheng et al. 2013).

Although five of the PS genes have no assigned function (Appendix, Table S3), OsTN2g002903 and OsTN1g003572 were identified as a PLATZ transcription factor (TF) family protein and an armadillo/beta-catenin repeat protein-like, respectively. Previous studies have shown that PLATZ TF GL6 in rice affects grain size and number (Wang et al. 2019a). In maize, PLATZs were found to be involved in the interaction with the RNA III polymerase (RNAP III) (Wang et al. 2018a). Specifically, mutational studies done on the PLATZ TF floury3 gene in maize endosperm resulted in inefficient production of RNAs in endosperm (Li et al. 2017). The OsTN2g002903 gene of TN1 also was mutated (Appendix, Figure S10c), and the amino acid change could have been advantageous to the plant.

The OsTN1g003572 gene (an armadillo/beta-catenin protein gene) can be associated with root development (Coates et al. 2006), disease resistance (Zeng et al. 2004) and abiotic stress response (Sharma et al. 2014). There is the possibility that it could have a role like that of PHOR1 (photoperiod-responsive protein 1), which is homologous to the armadillo protein of Drosophila. In potato, it influences tuberization and has been linked to the GA signaling pathway (Amador et al. 2001). Speaking of GA, OsTN5g001087, another gene under PS, is GA 3-beta-hydroxylase. It is an important enzyme in the final step of GA biosynthesis that produces the GA1 bioactive compound (Reinecke et al. 2013). OsTN5g001087 has been tagged with GO:0009416 (response to light stimulus) and is supported by the fact that GA biosynthesis is affected by light conditions (García-Martinez and Gil 2001). The alignment of GA 3β-hydroxylase proteins (Appendix, Figure S10d) shows that IR8 and MH63 are very similar to each other and the one to have changed amino acids at the PS sites was TN1.

Another PS gene is OsTN5g000040 (Table 4). It is a hypothetical protein and its orthologue in Nipponbare is also a hypothetical protein. For OsTN5g002486 and OsTN12g001576, no names were given to them nor do they have any protein orthologue in Nipponbare, maize or wheat.

## Conclusions

Using the available genome sequences of TN1 and IR8, we inferred that the current annotation of the semi-dwarf gene *sd1* contains errors. In particular, we found a 382-bp, instead of 383bp, genomic deletion, which resulted in a frameshift mutation. Sanger sequencing validated this deleted region in *sd1*, and we proposed a model of the *sd1* gene that corrects errors in the literature. We also predicted the blast disease resistant (R) genes of TN1 by finding TN1 orthologues of the R genes in Tetep, a well-known resistant cultivar. Haplotype analysis of the *Pi54* gene using cultivars from the 3,000 Rice Genomes Project revealed similar alleles of TN1 to a susceptible cultivar to blast, and different alleles when compared to resistant cultivars. In comparison, haplotype analysis of the *Pi-ta* gene of TN1 showed similar alleles to both resistant and susceptible cultivars. In addition, protein alignment of TN1 Pi54 against the blast resistant Tetep showed a loss of the first 598 amino acids. Of note, we found that *Pi54*, a well-known R gene, is absent in TN1, which partially explains why TN1 is more susceptible to blast than Tetep. We also scanned the TN1 genome using PosiGene, which is a software for detecting positively selected genes, and identified 11 genes deemed to have undergone positive selection in the past. Some of them are associated with drought-resistance and stress response. Our study fills some knowledge gaps in Green Revolution and in the study of the first semidwarf rice cultivar.

## Methods

Comparison of the *sd1* nucleotide and protein sequences of TN1, IR8 and Nipponbare Using the blast2go gff files of the TN1 and IR8 cultivars (TN1_blast2go_gff.gff and IR8_blast2go_gff.gff) available at https://figshare.com/articles/dataset/Green_Revolution_rice_genomes_annotation_fil es/13010333, the transcript ids corresponding to the *sd1* gene of TN1 and IR8 were obtained by the command: grep ’Gene=*sd1*’ <INPUT blast2go gff file>. The results were OsTN1t004133.1 and OsIR8_01T0407900.1 for TN1 and IR8, respectively. To get the corresponding gene ids, the suffix .1 in the transcript ids were dropped and the t or T was replaced by a g or a G. So, the *sd1* gene ids in TN1 and IR8 were OsTN1g004133 and OsIR8_01G0407900, respectively. For Nipponbare, querying “*sd1*” in the RAP-DB (Sakai et al. 2013) website gave Os01g0883800 as the gene id, while Os01t0883800-02 as its representative transcript, which has “GA 20-oxidase2, GA metabolism” as the description. GA stands for Gibberellin.

With gene ids now available, the nucleotide sequence corresponding to the gene region were obtained from their gff annotation file. Samtools (Li et al. 2009), version 1.8, extracted the gene sequence with this command: samtools faidx <INPUT genome fasta> <chromosome:start-end> > <OUTPUT gene fasta>. Gffread (Pertea and Pertea 2020), version 0.11.8, via default options, extracted the CDS and protein sequences of TN1, and samtools faidx command was executed to get the *sd1* CDS and protein of TN1. For IR8 and Nipponbare, because their CDS (oryza_indicair8.cds.fasta.gz and IRGSP-1.0_cds_2020-03-24.fasta.gz) and protein sequences (oryza_indicair8.cds.fasta.gz and IRGSP-1.0_protein_2020-03- 24.fasta.gz) were downloaded directly from their online repository (Appendix, Table S4), samtools faidx command was used to extract the sequences.

For the alignment involving nucleotide sequences, Clustal (Larkin et al. 2007), version 2.1 (parameter: -type=dna -align), was used. Inputs in Clustal were the concatenated fasta files. To create Figure 3, the Nipponbare CDS was used as input in https://www.bioline.com/media/calculator/01_13.html with one-letter translation selected as the output mode. The mapped amino acids to the respective codons of Nipponbare were checked against the CDS, protein sequences, as well as the boundaries of the exon regions of TN1 and IR8. Any such differences were indicated in Figure 3.

To know the deletion length in the TN1 gene with respect to the Nipponbare *sd1*, the Clustal alignment file was viewed in Jalview (Waterhouse et al. 2009), version 2.11.1.4. The same was also done for the alignment of the *sd1* genes between Nipponbare and IR8.

To create the gene structure models, GenePainter (Hammesfahr et al. 2013) was used. The input were the protein sequences of the genes (protein ids as fasta headers) aligned by Clustal (version 2.1, parameter: -output=FASTA -type=protein – align). The alignment file was uploaded to https://genepainter.motorprotein.de/genepainter, together with the segment of the gff annotation file containing the lines with the gene ids and transcript ids of the *sd1* of Nipponbare, TN1, and IR8. For the Nipponbare vs TN1 gene structure models, two gff files were prepared and named as Os01t0883800-02.gff and OsTN1t004133.1.gff, the filenames matching the fasta headers in the alignment file. The same method was done for the Nipponbare vs IR8.

### Sanger sequencing of the *sd1* gene

Genomic DNA was extracted from young leaves of TN1 and IR8 with DNeasy Plant Mini Kit (Qiagen). Primers for amplifying *sd1* gene were designed at the flanking regions about 150 bp upstream or downstream the target region. TN1 *sd1* gene was amplified by forward primer (5’-ATGTCTGTCCAGTGGCAACC-3’) and reverse primer (5’-CTTGAATTACTTGTTCTGTTGCTTC-3’) and IR8 *sd1* by forward primer (5’-ACCTTTAAACTTGGTCTAAAAGGATG-3’) and reverse primer (5’-GCTTGAATTACTTGTTCTGTTGC-3’) with ALLinTM Mega HiFi DNA polymerase (highQu). The result PCR products were purified with FB PCR Clean Up/ Gel Extraction Kit (Fair Biotech) and then sequenced by DNA Sequencing Core Facility of the Institute of Biomedical Sciences, Academia Sinica.

### Comparison of the *sd1* gene sequence against TN1 reads from the 3000 Rice Genomes Project

We first searched the SNP-Seek database (Mansueto et al. 2017) for any entry about the Taichung Native 1 cultivar by checking each results page and searching the page for key words like “Taichung” or “TN1”. We found two and they have the assay ids CX270 and CX162. The former has the entry “TAICHUNGNATIVE1”, while the latter is named as “TN1”. Alternatively, the search can be faster by doing this clicks in the SNP-Seek website: Home -> Download -> SNPs Analysis Files -> Variety drop down menu. We downloaded the Sequence Read Archive (SRA) reads associated with these entries in SNP-Seek and mapped them into the TN1 genome and checked their mapping coverage via (IGV) (Robinson et al. 2011).

To download the SRA reads, we use fastq-dump of the SRA Toolkit (SRA Tools 2021) version v2.10.5 (command: fastq-dump --split-files <SRA Accession ID>) to retrieve them as paired-end reads. We trimmed the reads using Trimmomatic (Bolger et al. 2014) v0.39 (parameters: adapters.fa:2:30:10 LEADING:20 TRAILING:20 SLIDINGWINDOW:4:20 MINLEN:50 CROP:82). The trimmed paired-end reads were interleaved by BBTools (Bushnell 2021) v38.90 (command: reformat.sh in1=read1.fastq in2=read2.fastq out=out.fastq). The output reads were mapped via BWA (Li 2013) (version 0.7.17) mem (default options) into the TN1 genome. Each output sam file was converted into a bam file via Samtools (Li et al. 2009) version 1.9 (commands: samtools view -S -h -b -f 3 -F 12 -q 20; samtools sort -T tmp; samtools index), then the filtered bam files were combined as one file with the merge command (parameter: -f -h <SAM file> --output-fmt BAM). The output bam file was sorted (command: samtools sort -T tmp) and indexed (default options). To get the mapping in the *sd1* region, the bam file was sliced (command: samtools view -b <INPUT bam file> ’TN1_chr1:40,361,934-40,364,270’). The mentioned coordinates in chromosome 1 of TN1 represent the locus of the *sd1* gene. The bam file was viewed in IGV (Eddy 1998) v2.10.0 through the Files tab and choosing Load from File. After loading the bam file, the TN1 genome fasta file was read in IGV through the Genomes tab and selecting Load Genome from File. Finally, the coordinates of the *sd1* gene locus (’TN1_chr1:40,361,934-40,364,270) was inputted in the search box and Go was clicked.

To retrieve the SNP data of the *sd1* gene, the Genotypes icon was clicked from the homepage of the SNP-Seek website. Os01g0883800 (RAPDB-ID of the *sd1* gene) was inputted in the Gene locus section. A dropdown menu appeared and Os01g0883800 was selected and automatically the CHR1, 38382385, and 38385469 became the values for Chromosome, Start and End, respectively. Variety set was “3k” and the SNP set was “3kfiltered”. In the options settings, Include Indels was checked. All other settings were default. The Search button was clicked. In the results table, the Subpopulation column title was clicked.

### Prediction of NBS-LRR genes

To detect the NBS-LRR genes of TN1, the proteins extracted from the gff annotation file and genome of TN1, via gffread (Pertea and Pertea 2020) (default options) were used as input in hmmscan (Eddy 1998), (via HMMER, version 3.1b2) and NLR-parser (Steuernagel et al. 2015), version 1.0 (Figure S8). By using the Pfam 30.0 database (El-Gebali et al. 2018), domains of the TN1 proteins were predicted.

The domain table output contained the list of predicted functional domain of each protein with their identified locations.

NLR-parser (Steuernagel et al. 2015) looked for R genes in the TN1 proteins based on a pre-defined set of NLR domains, classifying them either as an NB-ARC, LRR or NB-LRR. NLR-Parser needs an xml file as input and this was produced by the mast (Bailey and Gribskov 1998) [MEME Suite (Bailey et al. 2009), version 4.9.1] tool. The resulting output xml file became the input for the NLR-Parser run. The results of the hmmscan and NLR-parser runs were integrated and saved as TN1_genome.maker.pass2.maker_proteins.rename.NLRs.csv.

### Search for Tetep NLR orthologues in TN1

Input data was prepared by extracting first the coding sequence (CDS) of TN1, followed by getting the CDS of NB-ARC domains. Finally, the protein sequences of the NB-ARC domains were finally obtained. The protein fasta file served as the input file in the OrthoFinder (Emms and Kelly 2019) run.

To investigate why TN1 is susceptible to blast disease, the NB-ARC fasta files of Tetep were blasted to TN1 via OrthoFinder (Emms and Kelly 2019), version 2.2.7, blastp (Camacho et al. 2009) (NCBI BLAST+, version 2.3.0) and blastn (Camacho et al. 2009) (NCBI BLAST+, version 2.3.0) (Figure S9). OrthoFinder searched for orthologues of the TN1 NB-ARC domains against the NB-ARC domains of Tetep, Nipponbare, MH63 and R498. Results were organized and saved as TN1_Orthologues.ortho_pairs.csv. Identified orthologues between TN1 and Tetep were considered found with respect to TN1. If no orthologue was found, Tetep NB-ARC was aligned via blastp (Camacho et al. 2009) (parameter: -evalue 1e-10; output filename: Tetep_NBS_domain_protein.blastp.TN1_marker_protein.csv) against the set of TN1 proteins. The best results were saved as Tetep_NBS_domain_protein.blastp.TN1_marker_protein.best.csv. An R gene was found if it had more than 50% alignment coverage. The process was repeated for the NB-ARC of Tetep against MH63, R498 and Nipponbare.

The results of the mapping of the tested NLRs were saved as Tetep_NBS_domain_protein.blastp.TN1_marker_protein.best.tested.csv. The file Tetep_tested.csv was derived from Table S6 results of the Wang et al. (2019b) study and is available at https://doi.org/10.6084/m9.figshare.14546724. The file has two columns. Column 1 is Gene ID of the Tetep R gene. Column 2 follows this notation, Receptor:Total_Tested:Resistant:Susceptible. Receptor refers to either of TP309 or Shin2 as the receptor cultivars of the R gene. Total_Tested refers to the number of tested blast strains for the receptor cultivar, while Resistant and Susceptible are the counts of being resistant or susceptible to the pathogen. To confirm that whether those “absent” genes are deleted or possibly not coding anymore, we blasted Tetep NB-ARC domain CDS against TN1 genome via blastn. The results were organized and saved as Tetep_NBS_domain_cds.blastn.TN1_genome.best.csv.

To check the effect of sequence variation, the TN1 genome and the Tetep genome were aligned via MUMmer (Kurtz et al. 2004), version 3.23. Only unique alignments were used in variants calling. Effects of variants were predicted using snpEff (Cingolani et al. 2012), version 4.3o (parameter: -ud 2000, input: Tetep_v_TN1.nucmer1.filter.vcf.gz, output: Tetep_v_TN1.nucmer1.filter.snpEff.vcf): The output file was parsed by sum_snpEff.pl and map_records.pl script, and the results were saved as Tetep.NBS_genes.TN1_nucmer1.snpEff.csv.

The commands used in the search for Tetep NLR orthologues in TN1 are available at https://doi.org/10.6084/m9.figshare.14555598 as the Search_for_Tetep_NLR_orthologues_in_TN1_script.sh file. The code for sum_snpEff.v1.0.1.pl and for the prediction of NBS-LRR genes, and in the detection of presence or absence of blast R genes in TN1, is also available on the same Figshare link. The gff2fasta.pl, map_records.pl, and extract_split_seqs.pl scripts are available at https://github.com/wl13/BioScripts. For the mummer2Vcf.pl script, it is hosted at https://github.com/douglasgscofield/bioinfo/blob/master/scripts/.

A Chi-squared test was done using R (R Core Team 2021), to test whether the ratio of resistant/non-resistant NLR orthologues of TN1 were significantly different as compared to Tetep (Appendix, Table S2). R (version 3.6.1) command: chisq.test(c(69,170-69), p=c(90/219,1-90/219)). Result of the R command: X-squared = 0.018098, df = 1, p-value = 0.893.

### Detection of presence or absence of blast R genes in TN1

The study of Mahesh et al. (2016) was repeated for TN1 to detect whether a specific blast R gene was present or absent. Twenty-two cloned blast NLR protein sequences (Pib, *Pi-ta*, *Pi54*(Pik-h), Pid2, Pi9, Piz-t, Pi37, Pi36, Pik-m, pi21, Pit, Pi5, Pid3, Pb1, Pish, Pi25, Pia(RGA4), Pik-p, Pik, *Pi54*rh, Pi1, Pi64) were aligned via blastp (Camacho et al. 2009), version (NCBI BLAST+, version 2.3.0, parameter: -evalue 1e-10) to the TN1 protein sequences, and also by tblastn (Camacho et al. 2009), (NCBI BLAST+, version 2.3.0, parameter: -evalue 1e-10) against the TN1 genome to detect similar protein sequences. We get the hits which have an e-value < 10e-10 and identity >= 70%. The same method was applied to the Tetep proteins and genome sequence.

Missing R genes were denoted by a - sign and those that are found are given by a + mark, provided that the alignment sequences showed high similarity. An R gene was classified as mutated if there was a disagreement with the alignment, or the blastn best hit was better than the blastp result.

To find the Nipponbare orthologs of the TN1 blast R genes, OrthoFinder (Emms and Kelly 2019), version 2.3.11 (parameter: -S blast) was executed against the Nipponbare proteome from RAP-DB (Rice Annotation Project Database) (Sakai et al. 2013).

### Haplotype analysis using data from the 3,000 Rice Genomes Project

Haplotype analysis of the *Pi-ta* and *Pi54* genes were done in the SNP-Seek database (Mansueto et al. 2017) using the 3k filtered dataset. The objective is to get the haplotypes of the two genes. Starting from homepage of SNP-Seek, Genotype was clicked. Inputs in the Gene locus were the RAP-DB IDs of *Pi-ta* and *Pi54*. These were Os12g0281300 and Os11g0639100, respectively. In the options, Include Indels was also selected, while all other settings were default before executing the search. For *Pi-ta*, it resulted in a set of 3024 varieties with 42 SNP and 127 INDEL positions, while for *Pi54* it was 3024 varieties with 46 SNP and 24 INDEL positions. From the Table view of the results, the Haplotype tab was selected. The resulting haplotypes were regrouped using the autogroup and pamk options. Results about the variety order and grouping of the alleles were downloaded.

From the study of Jia et. al (2003), Wang et. al (2008) and Thakur et. al (2015), a list of resistant and susceptible cultivars to blast disease harboring the *Pi-ta* or *Pi54* gene were gathered. Each of the cultivars was checked to see whether the SNPs were listed in the SNP-Seek Database. To know whether they are in SNP-Seek, these series of clicks were done: Home -> Download -> SNPs Analysis Files. Another way is to check the variety order tab of Dataset S4. All possible combinations of naming the cultivar were tried for those containing numbers. For example, NANJING 11 was searched as NANJING11, NANJING-11 or NANJING 11. Keywords were also tried; e.g., for the cultivar PUSA BUSMATI 1, the query used was BASMATI and one of the hits was PUSA(BASMATI 1). The important/causal SNPs related to susceptibility (Jia et al. 2003; Wang et al. 2008; Thakur et al. 2015) were checked on the SNP effects data in Dataset S5 to find any similarity.

To build Table 2 and Table 3, the following series of steps were followed: 1) Get haplotypes of *Pi54* and *Pi-ta*; 2) find the cultivars from (Jia et al. 2003; Wang et al. 2008; Thakur et al. 2015) in SNP-Seek; 3) from the haplotypes, get the nucleotide position in which the SNPs are different (heterozygous) across all haplotype group; 4) list the heterozygous alleles for each cultivar; 5) list the number of mismatch SNPs per cultivar from the variety order tab of Dataset S4; 6) list the alleles in the heterozygous SNP positions for each haplotype group; 7) get the major and minor alleles and minimum allele frequency, from the graph portion of the tabular results of SNP-Seek, by clicking the line graph to find the right SNP position and see the information sought.

We were not able to find Tetep in the list of cultivars included in the 3K RGP so to get the SNPs of Tetep, its chromosome 11 (containing *Pi54*) and tig00012489 (containing *Pi-ta*) were aligned against their equivalent chromosomes in Nipponbare containing the said R genes. This was done via nucmer (default options) of the MUMmer version 4. The output delta file was used as an input in the show-snps (parameter: -C, default options) command. To get the alleles of Tetep, those corresponding to the coordinates of Nipponbare indicated in Table 2 and Table 3 were checked. If the coordinate was not found in the output show-snps, then the reference allele and the Tetep allele were assumed to be the same.

Clustal alignment was done for the *Pi54* and *Pi-ta* protein sequences of TN1 against Tetep (*Pi54*) and Yashiro-mochi (*Pi-ta*). The alignment file was viewed in Jalview (Waterhouse et al. 2009), version 2.11.1.4. Protein identifiers/GenBank accession numbers of the input protein sequences were: OsTN11t002257.1 for TN1 *Pi54*; chr11.fgenesh2107.1 for Tetep *Pi54*; OsTN12t001092.1 for TN1 *Pi-ta*; ACY25067.1 for Yashiro-mochi Pita. Creation of the images for the Clustal alignment were similar to the method done by Panibe (2021).

### Detection of genes subjected to positive selection in TN1

Coding sequences from 24 plant genomes (Appendix, Table S4) were used as input in PosiGene (Sahm et al. 2017), version 0.1 (parameters: -as=TN1 -rs=TN1 - ts=TN1 -nhsbr) to detect positive selection. Fifteen rice varieties or species were: five *indica* cultivars (TN1, IR8, MH63, IR64 and 9311), the Nipponbare reference genome, two wild species of the *indica* cultivar (*O. rufipogon* and *O. nivara*) plus seven non-Oryza sativa species (*O. barthii*, *O. brachyantha*, *O. glaberrima*, *O. glumipatula*, *O. punctata*, *O. meridionalis*, and *O. longistaminata*). Nine members of the grass family (*Brachypodium distachyon*, *Eragrostis tef*, *Leersia perrieri*, *Panicum hallii* fil2, *Panicum hallii* hal2, *Setaria italica*, *Sorghum bicolor*, *Triticum aestivumm*, *Zea mays*) were used as outgroups. This was to prevent the TN1 genome from becoming the last common ancestor in the species tree that PosiGene would create. The CDSs of TN1 and IR64 were extracted from their gff file via gffread (Pertea and Pertea 2020) (default options). The CDSs of the other cultivars were downloaded directly; see Appendix, Table S4.

Fasta headers were processed to follow an “isoform|gene” name format (ex. gene1.1|gene1) as required by PosiGene. This helped the software identify which isoforms were from the same gene. The tool was executed with TN1 as the as (anchor species) (most complete set of genes), rs (reference species) (basis for orthologue assignment), and ts (target species) (branch to test). This was to make sure that all the TN1 genes were tested for positive selection. The HomoloGene file for rice, which PosiGene recommends, was not used because it was based on Build 4.0 of Nipponbare, which is a *japonica* cultivar and outdated. The instructions in the PosiGene manual were followed to run the PosiGene.pl perl script. In the results output of PosiGene, those with FDR <0.05 are PS genes.

### PosiGene command

The PosiGene command below is for testing the branch leading to TN1 only: perl PosiGene.pl -o=TN1_GRgenes -as=TN1 -rs=TN1:folder/TN1_cds.fasta -tn=32 - ts=TN1 \

-nhsbr=TN1:folder/TN1_cds.fasta, \

IR64:folder/IR64_cds.fasta, IR8:folder/oryza_indicair8_cds.fasta, \

O_rufipogon:folder/oryza_rufipogon_cds.fasta, \

O_nivara:folder/Oryza_nivara_cds.fasta, \

MH63:folder/MH63_cds.fasta, \

Nipponbare:folder/IRGSP_cds.fasta,\

O_barthii:folder/Oryza_barthii_cds.fasta, \

O_brachyantha:folder/Oryza_brachyantha_cds.fasta, \

O_glaberrima:folder/Oryza_glaberrima_cds.fasta, \

O_glumipatula:folder/Oryza_glumipatula_cds.fasta, \

9311:folder/Oryza_*indica*_cds.fasta, \

O_longistaminata:folder/Oryza_longistaminata_cds.fasta, \

O_meridionalis:folder/Oryza_meridionalis_cds.fasta, \

O_punctata:folder/Oryza_punctata_cds.fasta, \

Brachypodium_distachyon:folder/Brachypodium_distachyon_cds.fasta, \

Eragrostis_tef:folder/Eragrostis_tef_cds.fasta,\

Leersia_perrieri:folder/Leersia_perrieri_cds.fasta, \

Panicum_hallii_fil2:folder/Panicum_hallii_fil2_cds.fasta, \

Panicum_hallii_hal2:folder/Panicum_hallii_hal2_cds.fasta, \

Setaria_italica:folder/Setaria_italica_cds.fasta, \

Sorghum_bicolor:folder/Sorghum_bicolor_cds.fasta, \

Triticum_aestivum:folder/Triticum_aestivum_cds.fasta, \

Zea_mays:folder/Zea_mays_cds.fasta

The IR8 genome was also scanned via PosiGene using the same set of CDSs to detect any PS genes in TN1 (parameters: -as=IR8 -rs=IR8 -ts=IR8 -nhsbr). Unfortunately, no IR8 gene got an FDR<0.05.

### REVIGO visualization of GO Terms

The GO terms of the TN1 genes under positive selection were visualized using the REVIGO (Supek et al. 2011) website (http://revigo.irb.hr/, accessed May 31, 2020). The inputs in REVIGO were the list of GO terms of all the proteins of the PS gene (Table S3). The online tool clustered the GO terms and selected the representative terms based on the cut-off value of similarity (also called dispensability), which is based on semantic distance computed by the SimRel algorithm. Settings for the PosiGene result: database with GO term sizes: whole UniProt; semantic similarity measure: SimRel; similarity cut-off value: 0.7.

Because the output is online, clicking the scatterplot will reveal the actual value of uniqueness when the user hovers their mouse pointer on a specific sphere. The tabular output listing all the inputted GO terms, their grouping as well as their corresponding frequency, uniqueness and dispensability values were downloaded from the website.

### Declarations

Ethics approval and consent to participate: Not applicable Consent for publication: Not applicable Availability of data and material: The genomes used in the study have the following GenBank accession numbers TN1 (GCA_018853525.1), IR8 (GCA_001889745.1), Tetep (GCA_004348155.2), MH63 (GCA_001623365.2), R498 (GCA_002151415.1), Nipponbare (GCA_001433935.1). The TN1 reads from the 3000 Rice Genome Sequencing Project assay CX270 have the SRA accession numbers: ERX576687, ERX576688, ERX576689, ERX576690, ERX576691, ERX576692, ERX576693, ERX576694, ERX576695, ERX576696, ERX576697, ERX576698, ERX576699, ERX576700. For assay CX162, the SRA accession numbers are ERX592032, ERX592033, ERX592034, ERX592035, ERX592036, ERX592037, ERX592038, ERX592039, ERX592040, ERX592041, ERX592042, ERX592043. Sources links of the coding sequences used in the PosiGene analysis are in Appendix, Table S4. The gff annotation file of IR8 is available at the Gramene, http://ftp.gramene.org/oge/release-3/gff3/oryza_indicair8/, while for R498 it is available at MBKBase, http://mbkbase.org/R498/. The Blast2GO annotation file of the genes of TN1 and IR8 are available at Figshare, https://doi.org/10.6084/m9.figshare.13010333. TN1’s gff annotation file can also be found at the Figshare link previously mentioned. The gff annotation file of Tetep is available at https://doi.org/10.6084/m9.figshare.7775810.v1. The datasets supporting the conclusions of this article are included within the article and its additional files.

## Authors’ contributions

Analysis of *sd1* genes, mapping of TN1 reads from the 3,000 Rice Genomes Project, genome-wide scan of genes under positive selection, REVIGO visualization, orthologue search between Nipponbare and TN1 proteins, haplotype analysis of *Pi54* and *Pi-ta*, by J.P.P.; prediction of NBS-LRR genes, search for Tetep NLR orthologues in TN1, detection of presence or absence of blast R genes in TN1, by L.W.; experiments including polymerase chain reaction amplification of TN1 and IR8 DNA for Sanger sequencing, by Y.-C.L.; advised the study, by C.-S.W.; designed, advised, and supervised the study, by W.-H.L. All authors participated in the preparation of the manuscript and agreed to submit this version of the manuscript.

## Acknowledgements

This study was supported by Academia Sinica (AS-TP-109-L10 and AS-KPQ-109-ITAR-TD05).

