## Supplementary material for "Identifying mutations in *sd1*, *Pi54* and *Pi-ta,* and positively selected genes of TN1, the first semidwarf rice in Green Revolution": TN1_Supplementary_Material

### I. SUPPLEMENTARY FIGURES

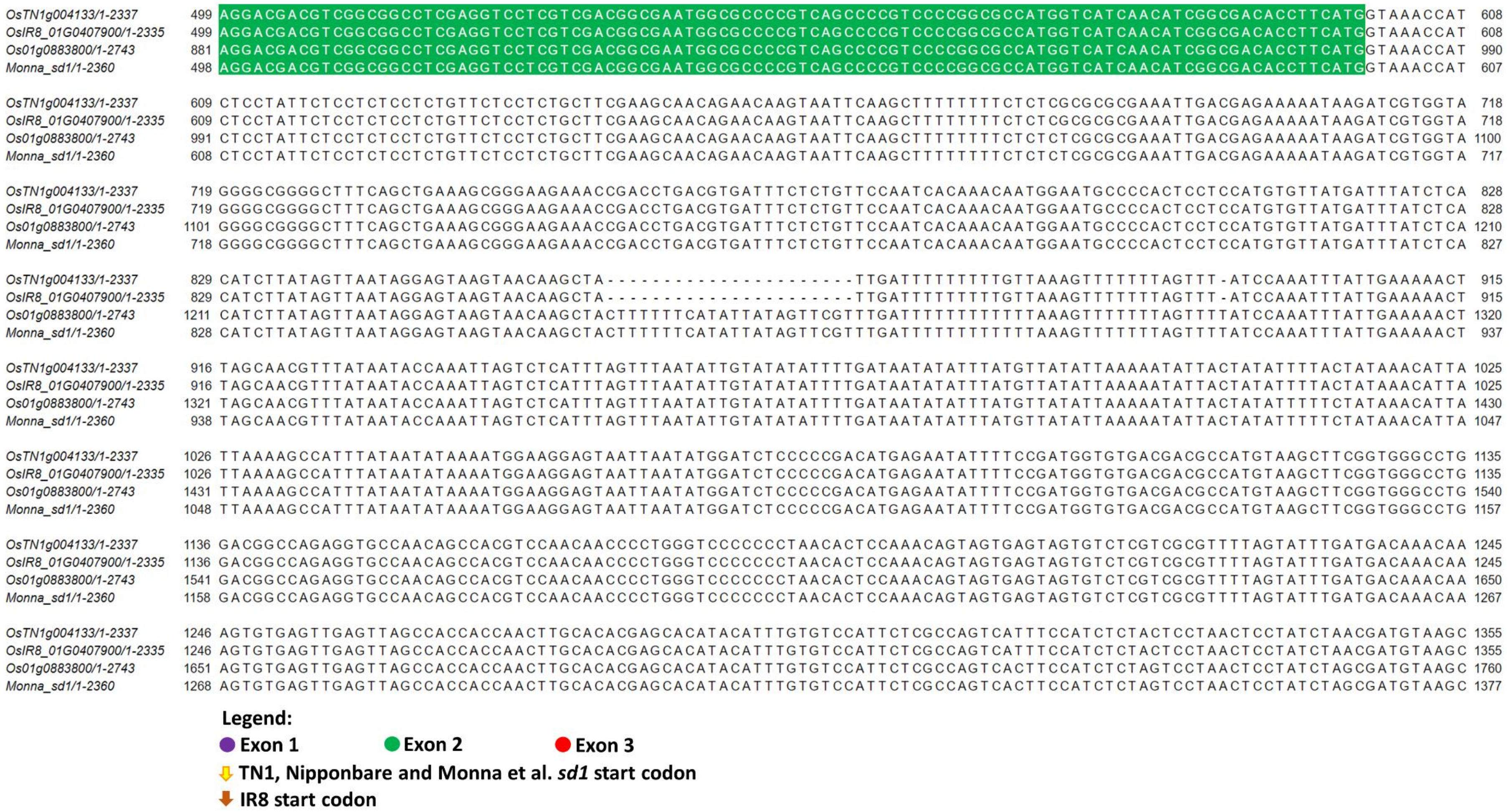


**Figure S1. Continuation of the *sd*1 gene-to-gene alignment in Figure 1**. The full gene sequence was used in the aligment which includes the untranslated region. Monna et al.'s *sd*1 sequence was copied from its publication. The alignment continues in the next image.

```
OsTN1g004133/1-2337        1356 GGATAATTTCATCATCCGTATATAAACCTGTTTGTTATAGTTAATTTCCTATATAATACTATAACAGTATACATTTTAAAAGAAAACAAAATTAGGATAAACAGGCCCTG 1465
OsIR8_01G0407900/1-2335    1356 GGATAATTTCATCATCCGTATATAAACCTGTTTGTTATAGTTAATTTCCTATATAATACTATAACAGTATACATTTTAAAAGAAAACAAAATTAGGATAAACAGGCCCTG 1465
Os01g0883800/1-2743        1761 GGATAATTTCATCATCCGTATATAAACCTGTTTGTTATAGTTAATTTCCTATATAATACTATAACAGTATACATTTTAAAAGAAAACAAAATTAGGATAAACAGGCCCTG 1870
Monna_sd1/1-2360           1378 GGATAATTTCATCATCCGTATATAAACCTGTTTGTTATAGTTAATTTCCTATATAATACTATAACAGTATACATTTTAAAAGAAAACAAAATTAGGATAAACAGGCCCTG 1487

OsTN1g004133/1-2337        1466 CTCCTATCCATCCATGGCACTTGGAAGGACCAGACTCGGTCATGCCATGCCAAGCCAAGATATGGATTATGGAAGAGTAGAGAAGAGGAGAGATGAGAGATAAGCATGCG 1575
OsIR8_01G0407900/1-2335    1466 CTCCTATCCATCCATGGCACTTGGAAGGACCAGACTCGGTCATGCCATGCCAAGCCAAGATATGGATTATGGAAGAGTAGAGAAGAGGAGAGATGAGAGATAAGCATGCG 1575
Os01g0883800/1-2743        1871 CTCCTATCCATCCATGGCACTTGGAAGGACCAGACTCGGTCATGCCATGCCAAGCCAAGATATGGGTTATGGAAGAGTAGAGAAGAGGAGAGATGAGAGATAAGCATGCG 1980
Monna_sd1/1-2360           1488 CTCCTATCCATCCATGGCACTTGGAAGGACCAGACTCGGTCATGCCATGCCAAGCCAAGATATGGGTTATGGAAGAGTAGAGAAGAGGAGAGATGAGAGATAAGCATGCG 1597

OsTN1g004133/1-2337        1576 TTCTCCTCCTCGTTGGATGTGTATTTTGGAGGGATTTGTGTAGTAGTAGCAGCGGCGCCGCGGGGACGGATGCGGATGGTGGCGCTTTCGGTGGCGTTTTCCTGGGGGG 1685
OsIR8_01G0407900/1-2335    1576 TTCTCCTCCTCGTTGGATGTGTATTTTGGAGGGATTTGTGTAGTAGTAGCAGCGGCGCCGCGGGGACGGATGCGGATGGTGGCGCTTTCGGTGGCGTTTTCCTGGGGG- 1684
Os01g0883800/1-2743        1981 TTCTCCTCCTCGTTGGATGTGTATTTTGGAGGGATTTGTGTAGTAGTAGCAGCGGCGCCGCGGGGACGGATGCGGATGGTGGCGCTTTCGGTGGCGTTTTCCCGGGGGGG 2090
Monna_sd1/1-2360           1598 TTCTCCTCCTCGTTGGATGTGTATTTTGGAGGGATTTGTGTAGTAGTAGCAGCGGCGCCGCGGGGACGGATGCGGATGGTGGCGCTTTCGGTGGCGTTTTCCCGGGGGGG 1707

OsTN1g004133/1-2337        1686 TTTTGGTTTGGCGCTTGGGGGGGATGGCATGGCGCGGCGTGCGGCTGCACGCCACACAC--GCGCGCGCACGCACGTACGTCGTCGTCGCCGCGGGCGGACGGTAGCTTA 1793
OsIR8_01G0407900/1-2335    1685 TTTTGGTTTGGCGCTTGGGGGG-ATGGCATGGCGCGGCGTGCGGCTGCACGCCACACAC--GCGCGCGCACGCACGTACGTCGTCGTCGCCGCGGGCGGACGGTAGCTTA 1791
Os01g0883800/1-2743        2091 TTTTGGTTTGGCGCTTGGGGGGGATGGCATGGCGCGGCGTGCGGCTGCACGCCACACACACGCGCGCGCACGCACGTACGTCGTCGTCGCCGCGGGCGGACGGTAGCTTA 2200
Monna_sd1/1-2360           1708 TTTTGGTTTGGCGCTTGGGGGGGATGGCATGGCGCGGCGTGCGGCTGCACGCCACACACACGCGCGCGCACGCACGTACGTCGTCGTCGCCGCGGGCGGACGGTAGCTTA 1817

OsTN1g004133/1-2337        1794 GGGTGGTGTGTTCCGCGCGCGGGCGCGGATTGTTCCATGCCGATCGATTTGGCGCCACCCTCGCCGCGGCTCTTGTCGCGTCGTGCGCCTCTCTCGCGCGGTTTGTCCTT 1903
OsIR8_01G0407900/1-2335    1792 GGGTGGTGTGTTCCGCGCGCGGGCGCGGATTGTTCCATGCCGATCGATTTGGCGCCACCCTCGCCGCGGCTCTTGTCGCGTCGTGCGCCTCTCTCGCGCGGTTTGTCCTT 1901
Os01g0883800/1-2743        2201 GGGTGGTGTGTTCCGCGCGCGGGCGCGGATTGTTCCATGCCGATCGATTTGGCGCCACCCTCGCCGCGGCTCTTGTCGCGTCGTGCGCCTCTCTCGCGCGGTTTGTCCTT 2310
Monna_sd1/1-2360           1818 GGGTGGTGTGTTCCGCGCGCGGGCGCGGATTGTTCCATGCCGATCGATTTGGCGCCACCCTCGCCGCGGCTCTTGTCGCGTCGTGCGCCTCTCTCGCGCGGTTTGTCCTT 1927

OsTN1g004133/1-2337        1904 GTCGCGTTGCTCAGCCGGCGACGGGGGCACGGACATTGGCGATGTAGCCCTGCACGTGTCGGCCTCTCCGTTGATGAATGATGATGTATGTATGTATTTTTTTTTTGTCT 2013
OsIR8_01G0407900/1-2335    1902 GTCGCGTTGCTCAGCCGGCGACGGGGGCACGGACATTGGCGATGTAGCCCTGCACGTGTCGGCCTCTCCGTTGATGAATGATGATGTATGTATGTATTTTTTTTTTGTCT 2011
Os01g0883800/1-2743        2311 GTCGCGTTGCTCAGCCGGCGACGGGGGCACGGACATTGGCGATGTAGCCCTGCACGTGTCGGCCTCTCCGTTGATGAATGATGATGTATGTATGTATTTTTTTTT-GTCT 2419
Monna_sd1/1-2360           1928 GTCGCGTTGCTCAGCCGGCGACGGGGGCACGGACATTGGCGATGTAGCCCTGCACGTGTCGGCCTCTCCGTTGATGAATGATGATGTATGTATGTATTTTTTTTT-GTCT 2036

OsTN1g004133/1-2337        2014 GAAGGAATTTGTGGGGAATTGTTGTGTGTGCAGGCGCTGTCGAACGGGAGGTATAAGAGCTGCCTGCACAGGGCGGTGGTGAACCAGCGGCGGGAGCGGCGGTCGCTGGC 2123
OsIR8_01G0407900/1-2335    2012 GAAGGAATTTGTGGGGAATTGTTGTGTGTGCAGGCGCTGTCGAACGGGAGGTATAAGAGCTGCCTGCACAGGGCGGTGGTGAACCAGCGGCGGGAGCGGCGGTCGCTGGC 2121
Os01g0883800/1-2743        2420 GAAGGAATTTGTGGGGAATTGTTGTGTGTGCAGGCGCTGTCGAACGGGAGGTATAAGAGCTGCCTGCACAGGGCGGTGGTGAACCAGCGGCGGGAGCGGCGGTCGCTGGC 2529
Monna_sd1/1-2360           2037 GAAGGAATTTGTGGGGAATTGTTGTGTGTGCAGGCGCTGTCGAACGGGAGGTATAAGAGCTGCCTGCACAGGGCGGTGGTGAACCAGCGGCGGGAGCGGCGGTCGCTGGC 2146

OsTN1g004133/1-2337        2124 GTTCTTCCTGTGCCCGCGGGAGGACAGGGTGGTGCGGCCGCCGCCGAGCGCCGCCACGCCGCGGCACTACCCGGACTTCACCTGGGCCGACCTCATGCGCTTCACGCAGC 2233
OsIR8_01G0407900/1-2335    2122 GTTCTTCCTGTGCCCGCGGGAGGACAGGGTGGTGCGGCCGCCGCCGAGCGCCGCCACGCCGCGGCACTACCCGGACTTCACCTGGGCCGACCTCATGCGCTTCACGCAGC 2231
Os01g0883800/1-2743        2530 GTTCTTCCTGTGCCCGCGGGAGGACAGGGTGGTGCGGCCGCCGCCGAGCGCCGCCACGCCGCAGCACTACCCGGACTTCACCTGGGCCGACCTCATGCGCTTCACGCAGC 2639
Monna_sd1/1-2360           2147 GTTCTTCCTGTGCCCGCGGGAGGACAGGGTGGTGCGGCCGCCGCCGAGCGCCGCCACGCCGCAGCACTACCCGGACTTCACCTGGGCCGACCTCATGCGCTTCACGCAGC 2256

OsTN1g004133/1-2337        2234 GCCACTACCGCGCCGACACCCGCACGCTCGACGCCTTCACGCGCTGGCTCGCGCCGCCGGCCGCCGACGCCGCCGCGACGGCGCAGGTCGAGGCGGCCAGCTGA 2337
OsIR8_01G0407900/1-2335    2232 GCCACTACCGCGCCGACACCCGCACGCTCGACGCCTTCACGCGCTGGCTCGCGCCGCCGGCCGCCGACGCCGCCGCGACGGCGCAGGTCGAGGCGGCCAGCTGA 2335
Os01g0883800/1-2743        2640 GCCACTACCGCGCCGACACCCGCACGCTCGACGCCTTCACGCGCTGGCTCGCGCCGCCGGCCGCCGACGCCGCCGCGACGGCGCAGGTCGAGGCGGCCAGCTGA 2743
Monna_sd1/1-2360           2257 GCCACTACCGCGCCGACACCCGCACGCTCGACGCCTTCACGCGCTGGCTCGCGCCGCCGGCCGCCGACGCCGCCGCGACGGCGCAGGTCGAGGCGGCCAGCTGA 2360
```

**Legend:**

● Exon 1 ● Exon 2 ● Exon 3

⇩ TN1, Nipponbare and Monna et al. *sd1* start codon

⬇ IR8 start codon

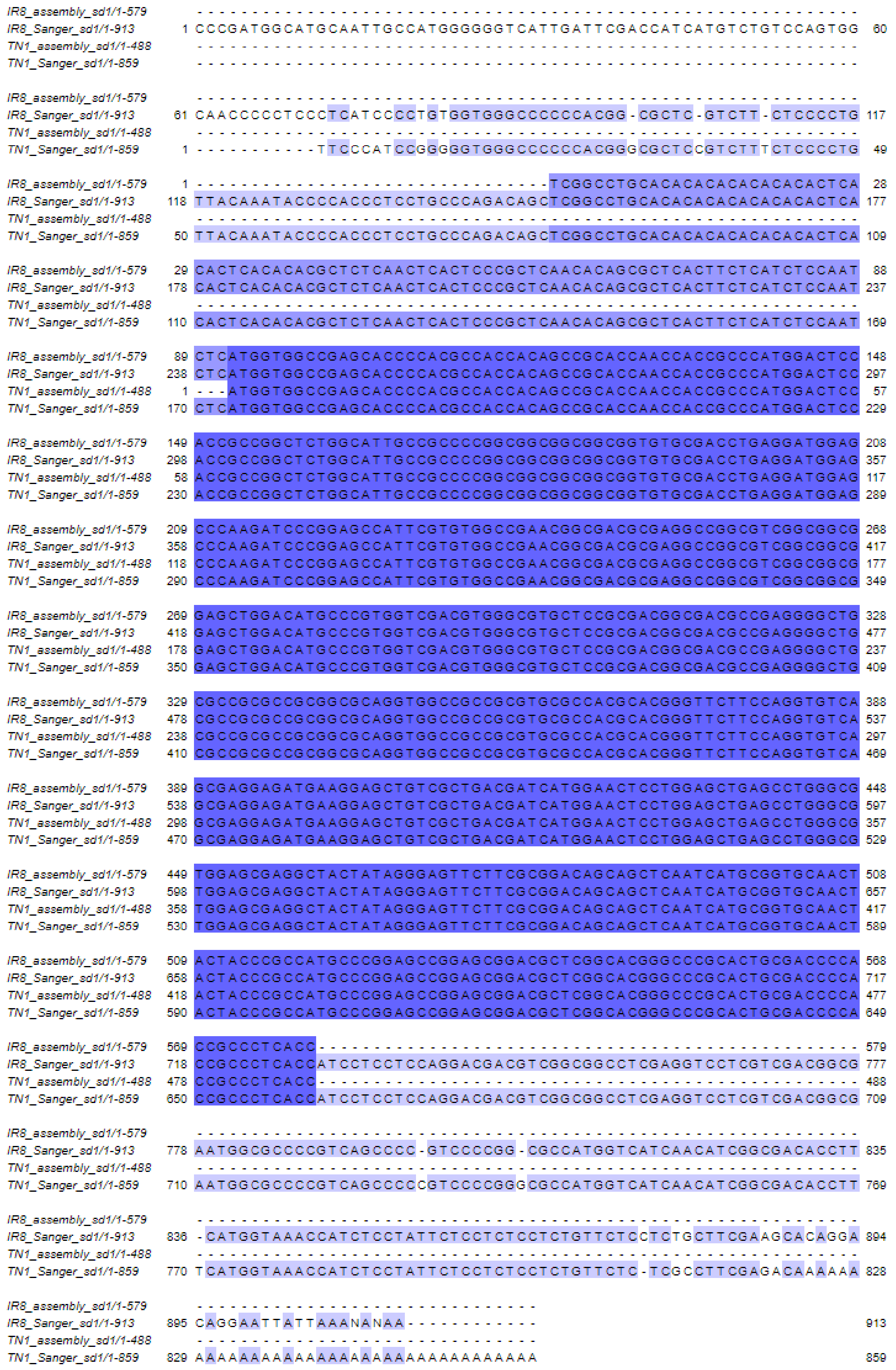


**Figure S2. Alignment of *sd*1 sequences of TN1 (chr 1:40,361,934-40,362,421), IR8 (chr 1: 39,824,196-39,824,774) vs. their Sanger sequencing.** Those highlighted in blue are 100% identical.

| Nipponbare positions | Variety | Assay | Accession | Subpopulation | Dataset | Mismatch | 38382846 | 38382847 | 38382848 | 38383054 | 38383055 | 38383056 | 38383057 | 38383058 | 38383059 | 38383060 | 38383061 | 38383062 | 38383063 | 38383064 | 38383065 | 38383066 | 38383221 | 38383286 | 38383461 | 38383462 | 38383463 | 38383464 | 38383465 |
|---|---|---|---|---|---|---|---|---|---|---|---|---|---|---|---|---|---|---|---|---|---|---|---|---|---|---|---|---|---|
| Nipponbare | | | | ▲ | | | C | C | G | T | T | C | A | A | A | T | T | C | A | A | A | A | C | C | A | T | T | C | T |
| VIETNAM ZAODAO | VIETI | B009 | IRGC | ind1A | 3k | 16.0 | C | — | — | T | T | C | A | A | A | T | T | C | A | A | A | A | T | C | A | T | T | C | T |
| KAHAMU | KAHA | B015 | IRGC | ind1A | 3k | 15.0 | C | — | — | T | T | C | A | A | A | T | T | C | A | A | A | A | T | C | A | T | T | C | T |
| TAIZHONGXIANXUAN 2 | TAIZH | B197 | | ind1A | 3k | 15.0 | C | — | — | T | T | C | A | A | A | T | T | C | A | A | A | A | T | C | A | T | T | C | T |
| TAICHUNGNATIVE1 | TAICI | CX27 | | ind1A | 3k | 14.5 | C | — | — | T | T | C | A | A | A | T | T | C | A | A | A | A | T | C | A | T | T | C | T |
| LAI YIP ZIM::IRGC 4955-1 | LAI Y | IRIS | IRGC | ind1A | 3k | 14.0 | C | C | G | T | T | C | A | A | A | T | T | C | A | A | A | A | T | C | A | T | T | C | T |
| CHI SHENG TAO::IRGC 4606-1 | CHI S | IRIS | IRGC | ind1A | 3k | 14.0 | C | C | G | T | T | C | A | A | A | T | T | C | A | A | A | A | T | C | A | T | T | C | T |
| SAN SHIH TSI::IRGC 1038-1 | SAN | IRIS | IRGC | ind1A | 3k | 14.0 | C | C | G | T | T | C | A | A | A | T | T | C | A | A | A | A | T | C | A | T | T | C | T |

**Figure S3**. **Screenshot of the genotype search result in the SNP-Seek database for TAICHUNGNATIVE1 (CX270).**

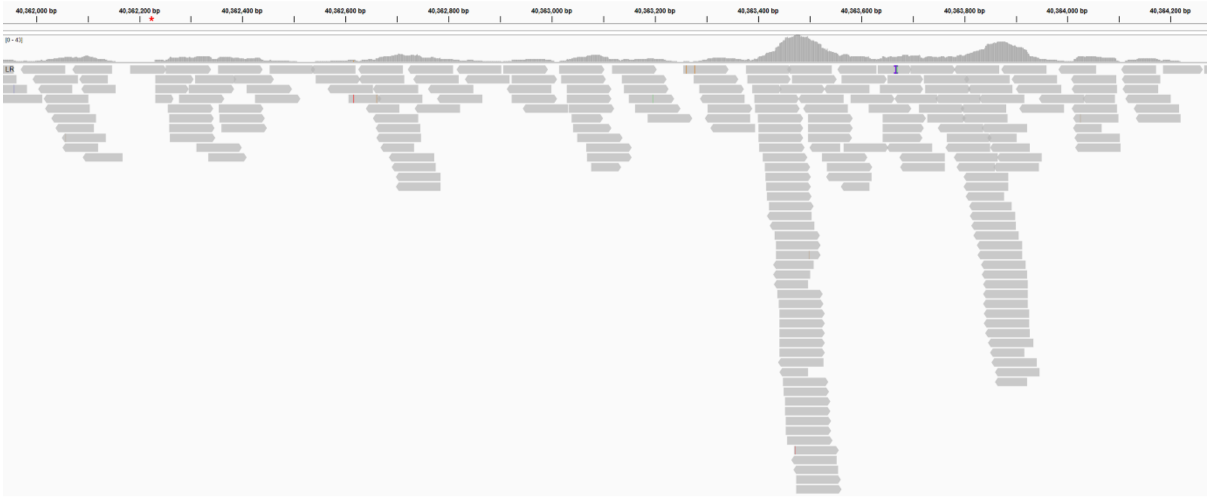


**Figure S4**. **Mapping of TAICHUNG NATIVE1 (CX270) properly paired reads in the TN1 sd1 gene region.** The red asterisk marks the location of the nucleotide that caused the 382-bp deletion in the sd1 gene of TN1 and IR8, instead of 383-bp.

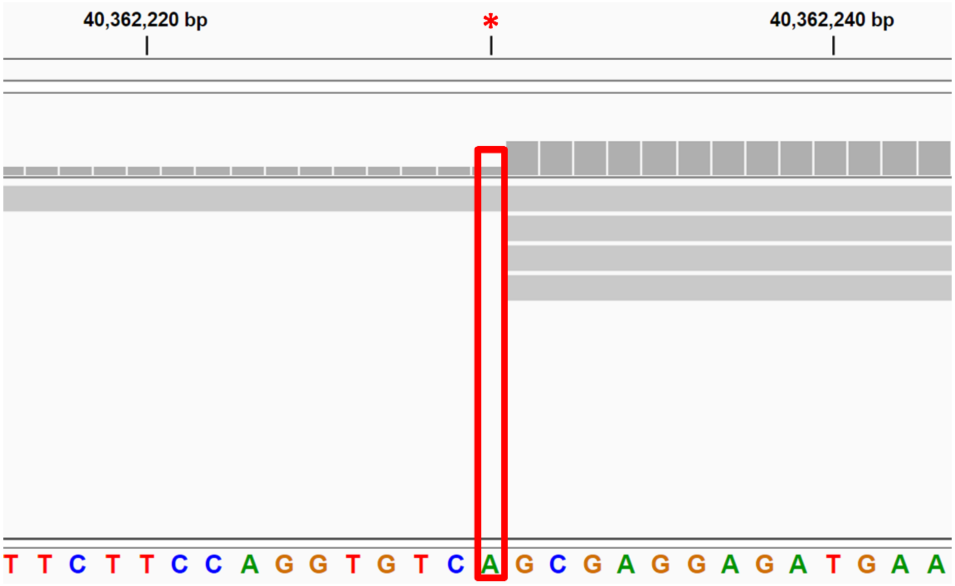


**Figure S5. Magnified view of the mapping of TAICHUNG NATIVE1 (CX270) reads at TN1 chromosome 1 position 40,362,230**. The letters are the nucleotides of TN1 chromosome 1. The red box indicates the region of interest. The vertical bars are the mapping coverage while the horizontal bars are the reads. The gray color of the reads means that their sequence is similar to the TN1 reference genome.

```
OsTN11t002257.1/1-337               ------------------------------------------------------------
chr11.fgenesh2107.1/1-1030       1  MEAAAGDLQAVMAMLSSSQMEEQCRLLRARSLLDEMRRAVADVHGALTATGATTSSVALA    60

OsTN11t002257.1/1-337               ------------------------------------------------------------
chr11.fgenesh2107.1/1-1030      61  ARTRSVCAPLRDALEAVLAMEGAATTNTLELSSLASAESSRSFLRYDSLEIERAVRANTI   120

OsTN11t002257.1/1-337               ------------------------------------------------------------
chr11.fgenesh2107.1/1-1030     121  RDALAEPMVGRSELAEKMVRVLLAVGGEEDGPLVMPIVGGPGIGKTRIVQALFNDSMVRE   180

OsTN11t002257.1/1-337               ------------------------------------------------------------
chr11.fgenesh2107.1/1-1030     181  KFPVRRWENVSERFNLFKMRMPNIWFNSTKFQNFLDEFINKSLNGRKGKYLVVLDDVWNE   240

OsTN11t002257.1/1-337               ------------------------------------------------------------
chr11.fgenesh2107.1/1-1030     241  NEAQDWPEWDSLMQALPSNGAVIFTTRTPMLVSKTAAVVPRTFPYFLQPLQQEHTVQFVH   300

OsTN11t002257.1/1-337               ------------------------------------------------------------
chr11.fgenesh2107.1/1-1030     301  QWLKRCWLDRSSEPFNIGMKIASKCDGVPLLIQSAGAILCRRPEAAFWQQFLEDFDVFFE   360

OsTN11t002257.1/1-337               ------------------------------------------------------------
chr11.fgenesh2107.1/1-1030     361  GSGLYSSDEEGSDILESAYSSYKHLPSHLQSCFLYCSMFPLGFNFDAEELADLFATAELT   420

OsTN11t002257.1/1-337               ------------------------------------------------------------
chr11.fgenesh2107.1/1-1030     421  GAQRIGFLEQLLNECFYPIEDSEYGGKFIYRMHKILHIFAVYMERELSTVVTADKDFTQV   480

OsTN11t002257.1/1-337               ------------------------------------------------------------
chr11.fgenesh2107.1/1-1030     481  QPSVRLMSLIIAPSTASFPRYIDQLKHLKALILLQDSRMLFSDQRCEIKEIDPMLCQSLK   540

OsTN11t002257.1/1-337            1  ----------------------------------------------------------LR     2
chr11.fgenesh2107.1/1-1030     541  HLQALSLQATKIRKLPNKIELVPHLRYLNLSQTNIETIPSSVSKLRLLQTLILSHCEKLW   600

OsTN11t002257.1/1-337            3  KLRENICKLVQLHKLDLEGCHYLVTLPTKKSKMKELQYLNVLNCYSLTAMPHAMGQLTHL    62
chr11.fgenesh2107.1/1-1030     601  KLHENICKLVQLHKLDLEGCLYLVTLPTKMSKMKKLQYLNVLNCYSLTAMPLAMGQLTNL   660

OsTN11t002257.1/1-337           63  HTLL--------------------------------------------------------    66
chr11.fgenesh2107.1/1-1030     661  HTLLGYFVPNNGSSAMSELQSLPDLNRLSLVNLEKVSDTEDARMAKLQEKEKLETLMLRW   720

OsTN11t002257.1/1-337           67  ----------DHEVLETLQPNQCLKTLEIVAYEGHAFPSWITSTEPYLTSLVEIRLVNLR   116
chr11.fgenesh2107.1/1-1030     721  NMDAGNASRIDHEVLETLQPSQCLKTLEIVAYEGYVFPSWMTRTEPYLTSLVEIRLVNMR   780

OsTN11t002257.1/1-337          117  SCEKRS----AIRTTDEVEISRVENISCIDDNFYGQNGTFPSLEKLILSFMTSLEVWEQS   172
chr11.fgenesh2107.1/1-1030     781  ACEKALPPLGILPCLKIAEISGVDNLSSIGDNFYGHNGTFPSLEKLILSYMTSLEVWEQS   840

OsTN11t002257.1/1-337          173  SRMNLFPRLAELVIIQCPKLRALHMEFPSIEKLILWMNNKMLYSSKEGLRGVEKSLENLS   232
chr11.fgenesh2107.1/1-1030     841  SRMNLFPRLAELVIIQCPKLRALHMEFPSIEKLILWMNNKMLYSSKEGMRGVEKSLENLS   900

OsTN11t002257.1/1-337          233  ISFCKELHASSGCEGLQALDRLKKLEICGCHELSCLPQGLQHLSSLASLKIDNCNKLEIL   292
chr11.fgenesh2107.1/1-1030     901  ISFCEELHASSGCEGLQALDRLKKLEICGCHELSCLPQGLQHLSSLTSLKIDNCNKLEIL   960

OsTN11t002257.1/1-337          293  PEWLENLPFLQIMCLSGCPILHSIPEGLTCSDIIVEYCPNFKEPS---------------   337
chr11.fgenesh2107.1/1-1030     961  PEWLENLPFLQIMCLSGCPILHSIPEGLTCSDIIVEDCPNFKEPSGMSSVLCSWKAMFLI  1020

OsTN11t002257.1/1-337               ----------
chr11.fgenesh2107.1/1-1030    1021  FIELFLKQLN                                                    1030
```

**Figure S6. Alignment of TN1 OsTN11t002257.1 against Tetep Pi54 (chr11.fgenesh2107.1).** Those highlighted in blue are 100% identical.

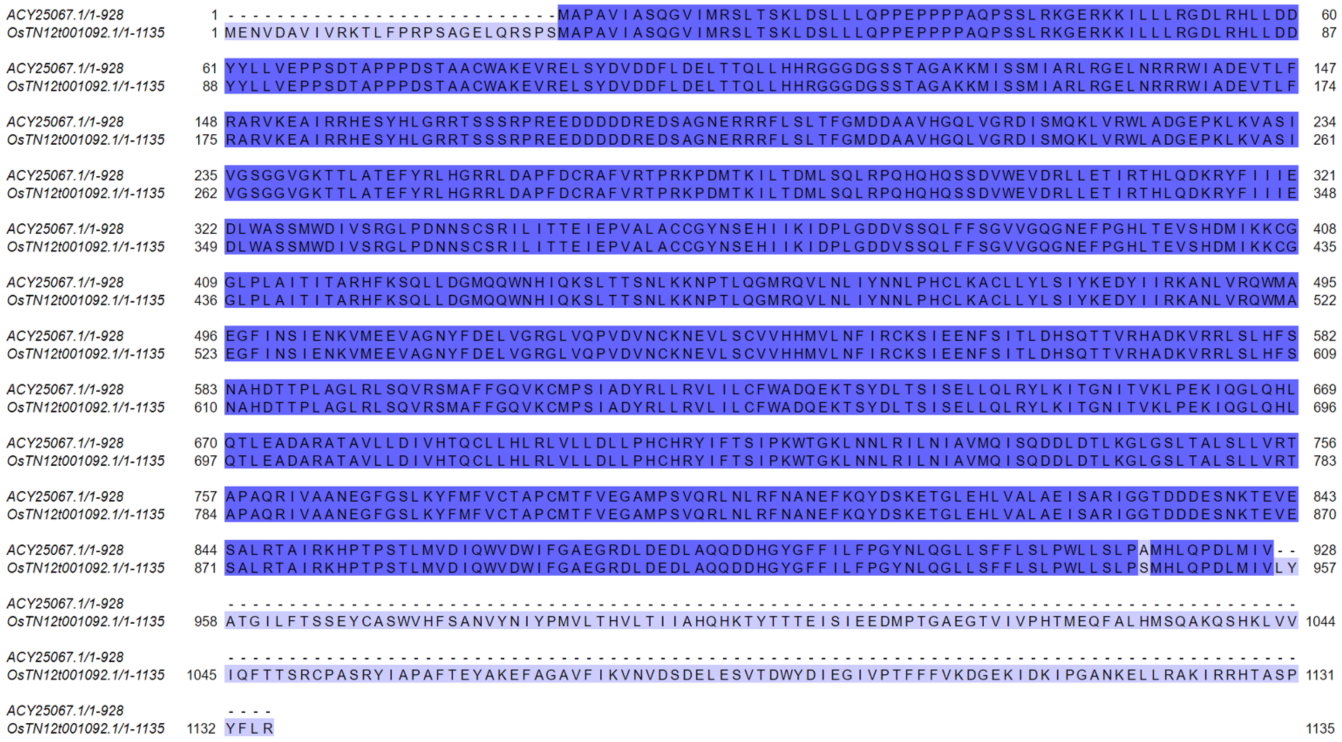


**Figure S7. Alignment of Pi-ta protein sequence of TN1 (OsTN12t001092.1) against Yashiro-mochi (ACY25067.1).** Those highlighted in blue are 100% identical.

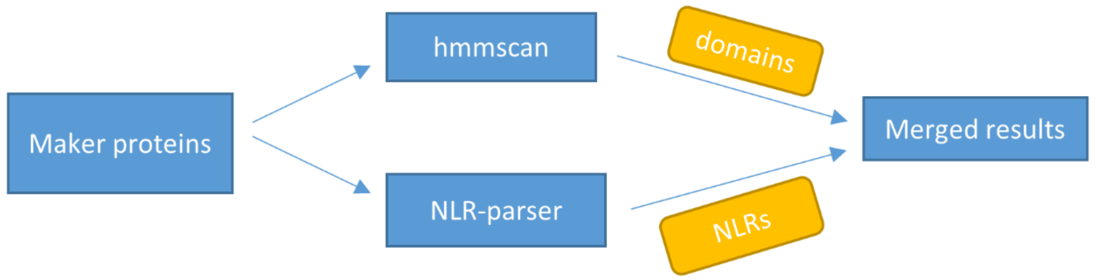


**Figure S8. R gene prediction workflow.** The merged data from hmmscan and NLR-parser represent the official results for the R gene search.

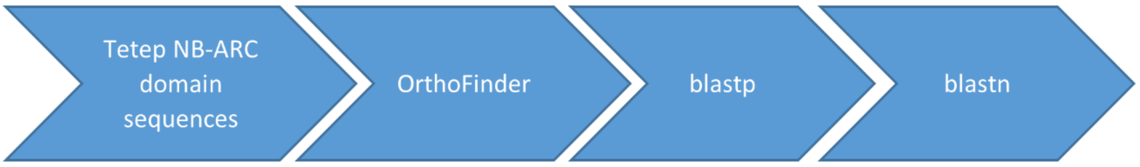


**Figure S9. Schematic diagram showing the Tetep R-gene search in the TN1 genome.** The inputs to OrthoFinder were protein sequences. For blastp, Tetep NB-ARC domains were searched against the TN1 proteins, while for blastn the Tetep NB-ARC coding sequences were aligned against the TN1 genome.

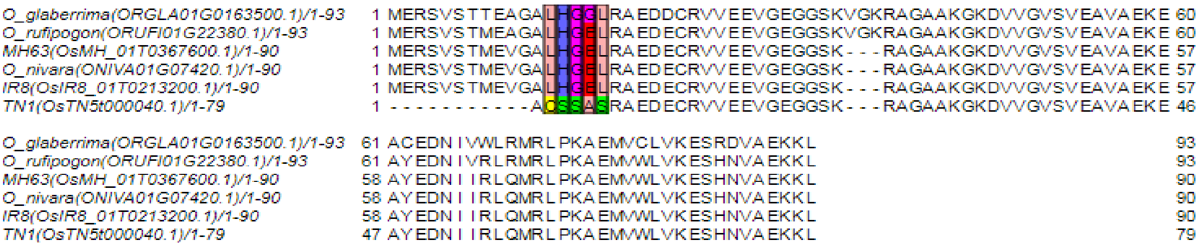


**Figure S10a. OsTN5g000040.1 (hypothetical protein) translated CDS alignment.** The TN1 PS sites and their probability of being under PS: Cys2 (99.45%), Ser3 (97.81%), Ser4 (98.05%), Ala5 (79.83%), and Ser6 (96.57%).

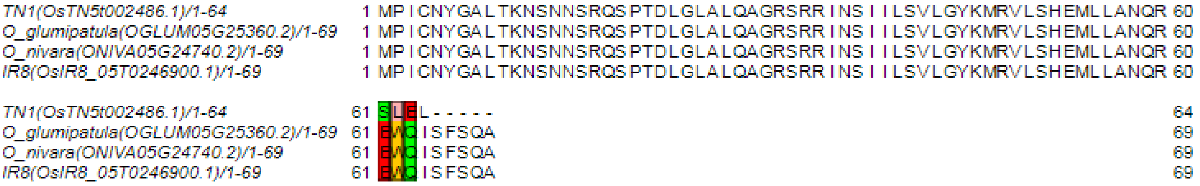


**Figure S10b. OsTN5g002486.1 translated CDS alignment.** The TN1 PS sites and their probability of being under PS: Ser61 (99.95%), Leu62 (99.01%), and Glu63 (99.02%).

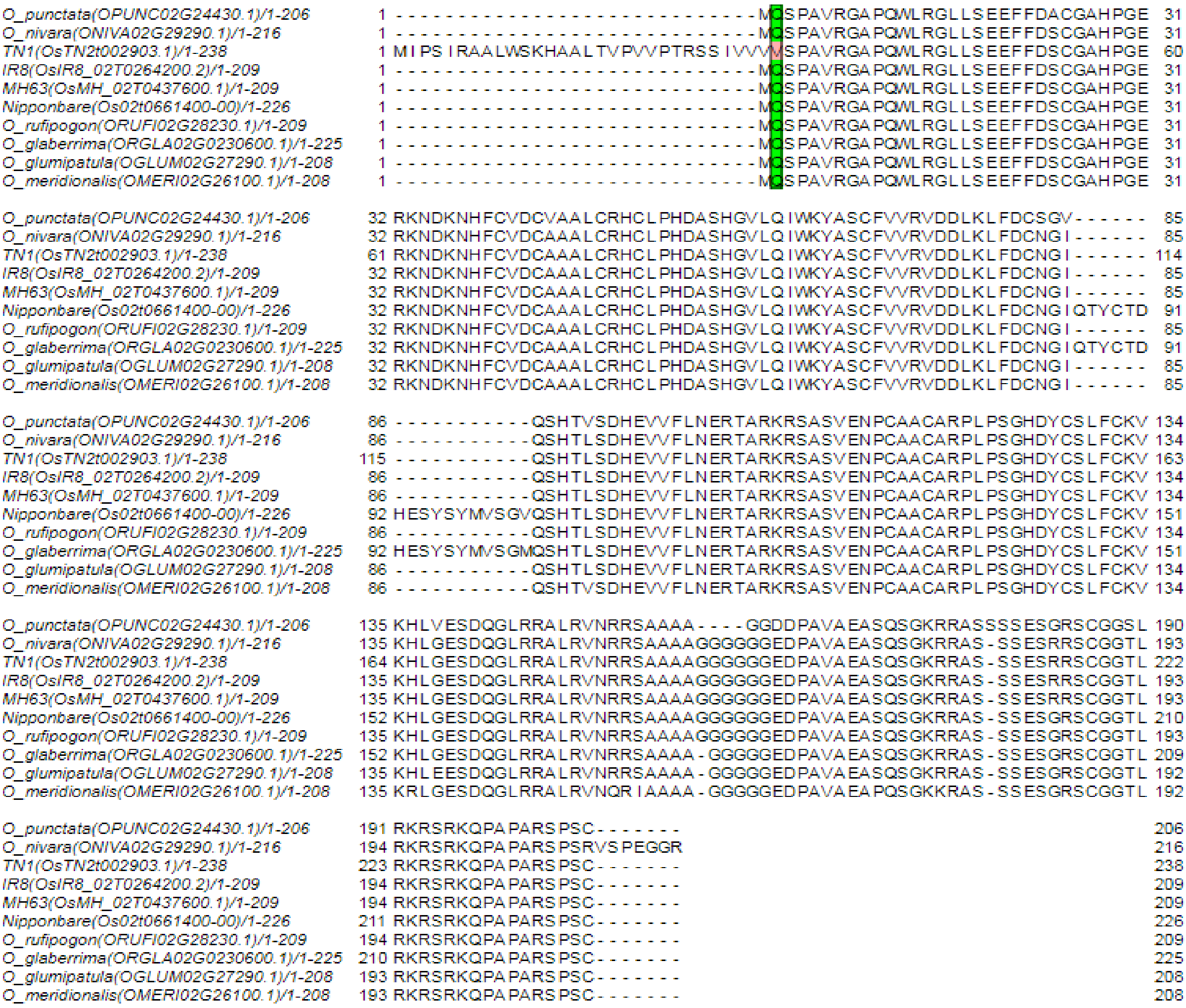


**Figure S10c. OsTN2g002903.1 (PLATZ transcription factor family protein) translated CDS alignment.** The TN1 PS site and the probability of being under PS: Val31 (99.33%).

```
O_nivara(ONIVA05G09470.1)/1-192          1 MEEFHKEIRRRRRAAGVVPQGRHWVSPESRWKKMTATMHWYPRCLDLRRALGLSAHSDSG  60
O_rufipogon(ORUFI05G09900.1)/1-192       1 MEEFHKEIRRRRRAAGVVPQGGHWVSPESRWKKMTATMHWYPRCPDLRRALGLSAHSNSG  60
O_glumipatula(OGLUM05G09430.1)/1-194     1 MEEFHKEMRRRRRAAGVVPQGGHWVSPESRWKKMTATMHWYPRCPDLRRALGLSAHSDSG  60
O_meridionalis(OMERI05G08150.1)/1-192    1 MEEFHKEMRRRRRAAGVVPQGGHWVSPESRWKKMTATMHWYPRCPDLRRALGLSAHSDSG  60
TN1(OsTN5t001087.1)/1-200                1 MEEFHKEIRRCRRAAGVVPQGGHWVSPESRWKKMTATMHWYPRCPDLRRALGLSAHSDSG  60
IR8(OsIR8_05T0099400.1)/1-180            1 MEEFHKEIRRCRRAAGVVPQGGHWVSPESRWKKMTATMHWYPRCPDLRRALGLSAHSDSG  60
MH63(OsMH_05T0143500.1)/1-180            1 MEEFHKEIRRCRRAAGVVPQGGHWVSPESRWKKMTATMHWYPRCPDLRRALGLSAHSDSG  60
O_barthii(OBART05G09240.1)/1-192         1 MEEFHKEIRRRRQAAGVVPQGGHWVSPESRWKKMTATMHWYPRCPDLRRALGLSAHSDSG  60

O_nivara(ONIVA05G09470.1)/1-192         61 FCSRASCQGCSYSGGDQTSGWQCRRSFLAPSSSTSATSSICMLTNGHFHSVYHRAVMNHN 120
O_rufipogon(ORUFI05G09900.1)/1-192      61 FCSRASCQGCSYSGGDQTGRWQCRRSFLAPSSSTSATSSICMLTNGRFHSVYHRAVMNRN 120
O_glumipatula(OGLUM05G09430.1)/1-194    61 FCSRASCQSCSYSGGDQTGGWQCRRSFLAPSSSTSATSSICMLTNGRFHSVYHRAVMNRN 120
O_meridionalis(OMERI05G08150.1)/1-192   61 FCSRASCQSCSYSGGDQTGGWQCRRSFLAPSSSTSATSSICMLTNGRFHSVYHRAVMNRN 120
TN1(OsTN5t001087.1)/1-200               61 FCSRASCQGCSCSGGDQTGGWQCRRSFLAPSSSTSATSSICMLTNGRFHSVYHRAVMNRN 120
IR8(OsIR8_05T0099400.1)/1-180           61 FCSRASCQGCSCSGGDQTGGWQCRRSFLAPSSSTSATSSICMLTNGRFHSVYHRAVFNVK 120
MH63(OsMH_05T0143500.1)/1-180           61 FCSRASCQGCSCSGGDQTGGWQCRRSFLAPSSSTSATSSICMLTNGRFHSVYHRAVFNVK 120
O_barthii(OBART05G09240.1)/1-192        61 FCSRASCQGCSCSGGDQTGGWQCRRSFLAPSSSTSATSSICMLTNGRFHSVYHRAVMNRN 120

O_nivara(ONIVA05G09470.1)/1-192        121 RDRISLGYFLGPLAER--RVPPGRSAAYRAVTWPEYKAVRKKAFTTGGSTLEMVSTPTAT 178
O_rufipogon(ORUFI05G09900.1)/1-192     121 RDRISLGYFLGPAAER--RVPPGRSAAYRAVTWPEYKAVRKKAFTTGGSTLEMVSTPTAT 178
O_glumipatula(OGLUM05G09430.1)/1-194   121 RDRISLGYFLGPPAGRQGRMPPGRSAAYRAVTWPEYKAVRKKAFTTGGSTLKMVSTPTAT 180
O_meridionalis(OMERI05G08150.1)/1-192  121 HDRISLGYFLGPPAGR--RMPPGRSAAYRAVTWPEYKAVRKKAFTTGGSTLKMVSTPTAT 178
TN1(OsTN5t001087.1)/1-200              121 RDRISLGYFLGLPAER--RVPPGWSAAYRAVTWPEYKAVRKKAFTTGGSTLEMVSTPTAT 178
IR8(OsIR8_05T0099400.1)/1-180          121 ------------VAGG--RVPPGWSAAYRAVTWPEYKAVRKKAFTTGGSTLEMVSTPTAT 166
MH63(OsMH_05T0143500.1)/1-180          121 ------------VAGG--RVPPGWSAAYRAVTWPEYKAVRKKAFTTGGSTLEMVSTPTAT 166
O_barthii(OBART05G09240.1)/1-192       121 RDRISLGYFLGPPAER--RVPPGRSAAYRAVTWPEYKAVRKKAFTTGGSTLEMVSTPTAT 178

O_nivara(ONIVA05G09470.1)/1-192        179 DEHNDVNK-------------IGKQYG                                  192
O_rufipogon(ORUFI05G09900.1)/1-192     179 DEHNDVADVVRDV--------I-----                                  192
O_glumipatula(OGLUM05G09430.1)/1-194   181 DEHNDVTDVVRDV--------I-----                                  194
O_meridionalis(OMERI05G08150.1)/1-192  179 DEHNDVTDVVQDV--------I-----                                  192
TN1(OsTN5t001087.1)/1-200              179 DEHNDVTDVVRDASILFERFGK-----                                  200
IR8(OsIR8_05T0099400.1)/1-180          167 DEHNDVTDVVRDV--------I-----                                  180
MH63(OsMH_05T0143500.1)/1-180          167 DEHNDVTDVVRDV--------I-----                                  180
O_barthii(OBART05G09240.1)/1-192       179 DEHNDVTDIVRDV--------I-----                                  192
```

**Figure S10d. OsTN5g001087.1 (GA 3β-hydroxylase) translated CDS alignment.** The TN1 PS sites and their probability of being under PS: Val117 (81.50%) and Arg119 (81.61%).

```
O_nivara(ONIVA01G38710.3)/1-275      1 MEVAECSEKLQFLKASASASMAYSIVQFPVKWQSIKYKLQKLCSNFQPEILNVPGDDGSC  60
TN1(OsTN1t003572.1)/1-221            1 MEVAECSEKLQFLKASASASMAYSIVQFPVKWQSIKYKLQKLCSNFQPEILNVPGDDGSC  60
IR8(OsIR8_01T0353800.1)/1-297        1 MEVAECSEKLQFLKASASASMAYSIVQFPVKWQSIKYKLQKLCSNFQPEILNVPGDDGSC  60
MH63(OsMH_01T0561700.1)/1-297        1 MEVAECSEKLQFLKASASASMAYSIVQFPVKWQSIKYKLQKLCSNFQPEILNVPGDDGSC  60
O_barthii(OBART01G33910.1)/1-257     1 MEVAECSEKLQFLKASASASMAYSIVQFPVKWQSIKYKLQKLCSNFQPEILNVPGDDDSC  60

O_nivara(ONIVA01G38710.3)/1-275     61 NEHVILVQFLQTAMATVSHIQAIASQCSDESYNGGRLPPPRARRSSPGARTHCPCPAAPP 120
TN1(OsTN1t003572.1)/1-221           61 NEHVILVQFLQTAMATVSHIQAIASQCSDESYNGGRLPPPRARRSSPGARTHCPCPAAPP 120
IR8(OsIR8_01T0353800.1)/1-297       61 NEHVILVQFLQTAMATVSHIQAIASQCSDESYNGGRLPPPRARRSSPGARTHCPCPAAPP 120
MH63(OsMH_01T0561700.1)/1-297       61 NEHVILVQFLQTAMATVSHIQAIASQCSDESYNGGRLPPPRARRSSPGARTHCPCPAAPP 120
O_barthii(OBART01G33910.1)/1-257    61 NEHVILVQFLQTAMATVSHIQAIASQCSDESYNGGRLPPPRARRSSPGARTHCPCPAAPP 120

O_nivara(ONIVA01G38710.3)/1-275    121 SAAARTQLLHLDATARSTPPRCGALLPRLRCRVARHRPPTFSSCPASHCQPPLHSLKSPA 180
TN1(OsTN1t003572.1)/1-221          121 SAAARTQLLHLDATARSTPPRCGALLPRLRCRVARHRPPTFSSCPASHCQPPLHSLKSPA 180
IR8(OsIR8_01T0353800.1)/1-297      121 SAAARTQLLHLDATARSTPPRCGALLPRLRCRVARHRPPTFSSCPASHCQPPLHSLKSPA 180
MH63(OsMH_01T0561700.1)/1-297      121 SAAARTQLLHLDATARSTPPRCGALLPRLRCRVARHRPPTFSSCPASHCQPPLHSLKSPA 180
O_barthii(OBART01G33910.1)/1-257   121 SAAARTQLLHLDATARSTPPRCGALLPRLRCRVARHRPPTFSSCPASHCQPPLHSLKSPA 180

O_nivara(ONIVA01G38710.3)/1-275    181 SSSFAACSYPSAARGAETSPLSPTGAQYSRRRAEAAPRRGRRETRVAERRWALA------ 234
TN1(OsTN1t003572.1)/1-221          181 SSSFAACSYPSAARGAETSPLSPTGAQYSRRRAEAAPRSDR------------------- 221
IR8(OsIR8_01T0353800.1)/1-297      181 SSSFAACSYPSAARGAETSPLSPTGAQYSRRRAEAAPRRGRRETRVAERRWALASWRNLH 240
MH63(OsMH_01T0561700.1)/1-297      181 SSSFAACSYPSAARGAETSPLSPTGAQYSRRRAEAAPRRGRRETRVAERRWALASWRNLH 240
O_barthii(OBART01G33910.1)/1-257   181 SSSFAACSYPSAARGAETSPLSPMGAN-SRRRAEAAPRRGRRETRVAERRWALASWRNLH 239

O_nivara(ONIVA01G38710.3)/1-275    235 ---------------------------------------------------------SCN 237
TN1(OsTN1t003572.1)/1-221              ------------------------------------------------------------
IR8(OsIR8_01T0353800.1)/1-297      241 KRGADMDKLTCGAHVDPALTQQQRWTKPVSKPPRDLCKRPGNGRMGWSPWELSSKWR--- 297
MH63(OsMH_01T0561700.1)/1-297      241 KRGADMDKLTCGAHVDPALTQQQRWTKPVSKPPRDLCKRPGNGRMGWSPWELSSKWR--- 297
O_barthii(OBART01G33910.1)/1-257   240 KRGADMDKLICGAHVIP---------------------------------------R--- 257

O_nivara(ONIVA01G38710.3)/1-275    238 GALTAAWGPQGEPRGAAPSSSSQTVRPRASGDAPDPTR                       275
TN1(OsTN1t003572.1)/1-221              --------------------------------------
IR8(OsIR8_01T0353800.1)/1-297          --------------------------------------
MH63(OsMH_01T0561700.1)/1-297          --------------------------------------
O_barthii(OBART01G33910.1)/1-257       --------------------------------------
```

**Figure S10e. OsTN1g003572.1 (armadillo/beta-catenin repeat protein-like) translated CDS alignment.** The TN1 PS sites and their probability of being under PS: Ser219 (85.68%) and Asp220 (79.28%).

```
O_punctata(OPUNC01G31310.2)/1-682   1 MTQTPSPAAAAPA---AEAGLPDAIAAALPPDPYEQLEVARKITAVAVAARASRLELEAAR  58
TN1(OsTN1t003413.1)/1-679           1 MTQTPTPAP-APAVAVSEAGLPDAIAAALPPDPYEQLEVARKITAVAVAARASRLELEAAR  60
MH63(OsMH_01T0541300.1)/1-623       1 MTQTPTPAP-APAVAVSEAGLPDAIAAALPPDPYEQLEVARKITAVAVAARASRLELEAAR  60
IR8(OsIR8_01T0339200.1)/1-623       1 MTQTPTPAP-APAVAVSEAGLPDAIAAALPPDPYEQLEVARKITAVAVAARASRLELEAAR  60

O_punctata(OPUNC01G31310.2)/1-682  59 LRQKLAEKDRLAAELADRTASLEQALRDSDARLCAALDDNAKLAKERDSLAHTSKKLARDL 119
TN1(OsTN1t003413.1)/1-679          61 LRQKLADKDRLAAELADRAASLEQALRDSDARLRAALDDNAKLAKERDSLAHTSKKLARDL 121
MH63(OsMH_01T0541300.1)/1-623      61 LRQKLADKDRLAAELADRAASLEQALRDSDARLRAALDDNAKLAKERDSLAHTSKKLARDL 121
IR8(OsIR8_01T0339200.1)/1-623      61 LRQKLADKDRLAAELADRAASLEQALRDSDARLRAALDDNAKLAKERDSLAHTSKKLARDL 121

O_punctata(OPUNC01G31310.2)/1-682 120 AKLETFKRHLMQSLGDDNPPIQETVDIRTCEQSVAKANSWK-------------------- 160
TN1(OsTN1t003413.1)/1-679         122 AKLETFKRHLMQSLGDDNPPIQETVDIRTCEQSVAKASSWKELHCYCTHIHTVHLFIRMHL 182
MH63(OsMH_01T0541300.1)/1-623     122 AKLETFKRHLMQSLGDDNPPIQETVDIRTCEQSVAKASSWK-------------------- 162
IR8(OsIR8_01T0339200.1)/1-623     122 AKLETFKRHLMQSLGDDNPPIQETVDIRTCEQSVAKASSWK-------------------- 162

O_punctata(OPUNC01G31310.2)/1-682 161 -----DGVAHSHHHHHHHPVSSLADGSTEIESV---------NQEVARPFEQKLTVTHITP 207
TN1(OsTN1t003413.1)/1-679         183 CNISEDGVAHSRHH---HPVSSLADGSTEIESLILLARVPAPNKTVARPFEQKLSVTHISP 240
MH63(OsMH_01T0541300.1)/1-623     163 -----DGVAHSRHH---HPVSSLADGSTEIESV---------NQEVARPFEQKLSVTHISP 206
IR8(OsIR8_01T0339200.1)/1-623     163 -----DGVAHSRHH---HPVSSLADGSTEIESV---------NQEVARPFEQKLSVTHISP 206

O_punctata(OPUNC01G31310.2)/1-682 208 RLTSDPAAKMRSAVTSPRRYSTAVSPKPAAAAASPRLEGHMAMQPWLPSSKMSSAANSPPR 268
TN1(OsTN1t003413.1)/1-679         241 RLTSDPAAKTRTAATSPRRYSTAVSPKLAASATSPRLEGHMAMQPWLLSSKMSSAANSPPR 301
MH63(OsMH_01T0541300.1)/1-623     207 RLTSDPAAKTRTAATSPRRYSTAVSPKLAASATSPRLEGHMAMQPWLLSSKMSSAANSPPR 267
IR8(OsIR8_01T0339200.1)/1-623     207 RLTSDPAAKTRTAATSPRRYSTAVSPKLAASATSPRLEGHMAMQPWLLSSKMSSAANSPPR 267

O_punctata(OPUNC01G31310.2)/1-682 269 GHSISGRTTRVDGKEFFRQARNRLSYEQFAAFLANIKELNAQRQSREETLQKADEIFGSEN 329
TN1(OsTN1t003413.1)/1-679         302 AHSISGRTTRVDGKEFFRQARNRLSYEQFAAFLANIKELNAHRQSREETLQKADEIFGSEN 362
MH63(OsMH_01T0541300.1)/1-623     268 AHSISGRTTRVDGKEFFRQARNRLSYEQFAAFLANIKELNAHRQSREETLQKADEIFGSEN 328
IR8(OsIR8_01T0339200.1)/1-623     268 AHSISGRTTRVDGKEFFRQARNRLSYEQFAAFLANIKELNAHRQSREETLQKADEIFGSEN 328

O_punctata(OPUNC01G31310.2)/1-682 330 KDLFMSFQGLLSP-----------------------GGTGGRPSLSRESATPPRHLASAAG 367
TN1(OsTN1t003413.1)/1-679         363 KDLFMSFQSLLSPLLIWTLHAVLQLDHERQITDGTA------------------------- 398
MH63(OsMH_01T0541300.1)/1-623     329 KDLFMSFQSLLSP------------------------------------------------ 341
IR8(OsIR8_01T0339200.1)/1-623     329 KDLFMSFQSLLSP------------------------------------------------ 341

O_punctata(OPUNC01G31310.2)/1-682 368 ERARRRAADPDLASSTGAAYSRGAGPASELRFVGGLCGRHPSTANLHLGRNRTE------- 421
TN1(OsTN1t003413.1)/1-679         399 ------------------------------------------------------VFACTET 405
MH63(OsMH_01T0541300.1)/1-623     342 ------------------------------------------------------VFACTET 348
IR8(OsIR8_01T0339200.1)/1-623     342 ------------------------------------------------------VFACTET 348

O_punctata(OPUNC01G31310.2)/1-682 422 --------------GVKVYQMMDLSTTSVVAAKAYKYKAESLVKDYLLADSYVSYTAVLGG 468
TN1(OsTN1t003413.1)/1-679         406 NNSLIMPRNATDRSGVKVYQMMDLSTTSVVAAKAYKYRAESLVKDYLLADCYVSYTAVLGG 466
MH63(OsMH_01T0541300.1)/1-623     349 NNSLIMPRNATDRSGVKVYQMMDLSTTSVVAAKAYKYRAESLVKDYLLADCYVSYTAVLGG 409
IR8(OsIR8_01T0339200.1)/1-623     349 NNSLIMPRNATDRSGVKVYQMMDLSTTSVVAAKAYKYRAESLVKDYLLADCYVSYTAVLGG 409

O_punctata(OPUNC01G31310.2)/1-682 469 ILMCKMVYDITHLISSLYYKGYGSLTKIQKLEWNNRGMSTVHAMFITLMSVYLVFFSNLFS 529
TN1(OsTN1t003413.1)/1-679         467 ILMCKMVYDITHLISSLYYKGYGSLTKIQKLEWNNRGMSTVHAMFITLMSVYLVFFSNLFS 527
MH63(OsMH_01T0541300.1)/1-623     410 ILMCKMVYDITHLISSLYYKGYGSLTKIQKLEWNNRGMSTVHAMFITLMSVYLVFFSNLFS 470
IR8(OsIR8_01T0339200.1)/1-623     410 ILMCKMVYDITHLISSLYYKGYGSLTKIQKLEWNNRGMSTVHAMFITLMSVYLVFFSNLFS 470

O_punctata(OPUNC01G31310.2)/1-682 530 DELDGPVTVRSSNLSNFTLGVSLGYFTADLAMIFWAYPSLGGMEYVLHHLLSIISLVYAIY 590
TN1(OsTN1t003413.1)/1-679         528 DELDGPVTVRSSNLSNFTLGVSLGYFIADLAMLSWAYPSLGGMEYVLHHLLSIISLVYAIY 588
MH63(OsMH_01T0541300.1)/1-623     471 DELDGPVTVRSSNLSNFTLGVSLGYFIADLAMLSWAYPSLGGMEYVLHHLLSIISLVYAIY 531
IR8(OsIR8_01T0339200.1)/1-623     471 DELDGPVTVRSSNLSNFTLGVSLGYFIADLAMLSWAYPSLGGMEYVLHHLLSIISLVYAIY 531

O_punctata(OPUNC01G31310.2)/1-682 591 SEEGQLYTYMVLISETTTPGINLRWFLDTVGMKRSKAYLVNGVTMFVAWLVKQMRTFSCIL 651
TN1(OsTN1t003413.1)/1-679         589 SEEGQLYTYMVLISETTTPGINLRWFLDTVGMKRSKAYLVNGVTMFVAWLVKQMRTFSCIL 649
MH63(OsMH_01T0541300.1)/1-623     532 SEEGQLYTYMVLISETTTPGINLRWFLDTVGMKRSKAYLVNGVTMFVAWLVKQMRTFSCIL 592
IR8(OsIR8_01T0339200.1)/1-623     532 SEEGQLYTYMVLISETTTPGINLRWFLDTVGMKRSKAYLVNGVTMFVAWLVKQMRTFSCIL 592

O_punctata(OPUNC01G31310.2)/1-682 652 IFAVPTILLVMNTVWFAKILRGLKKTLAKRQ                               682
TN1(OsTN1t003413.1)/1-679         650 IFAVPTILLVMNTVWFVKILRGLKKTLAKR-                               679
MH63(OsMH_01T0541300.1)/1-623     593 IFAVPTILLVMNTVWFVKILRGLKKTLAKRQ                               623
IR8(OsIR8_01T0339200.1)/1-623     593 IFAVPTILLVMNTVWFVKILRGLKKTLAKRQ                               623
```

**Figure S10f. OsTN1g003413.1 (transmembrane protein 56 isoform X1) translated CDS alignment.** The TN1 PS sites and their probability of being under PS: Lys223 (80.52%) and Thr224 (96.54%).

```
IR8(OsIR8_08T0127600.1)/1-194          1 ----------------------MSKGQMVHEIHSKNSWNKEMRLSTCSTALDSTLKTCSR  38
MH63(OsMH_08T0238500.1)/1-194          1 ----------------------MSKGQMVHEIHSKNSWNKEMRLSTCSTALDSTLKTCSR  38
O_nivara(ONIVA04G28960.1)/1-201        1 ----------------------MSKGQMVHEIHSKNSWNKEMRLSTCSTALDSTLKTCSR  38
O_rufipogon(ORUFI08G12790.1)/1-184     1 ----------------------MSKGQMVHEIHSKNSWNKEMRLSTCSTALDSTLKTCSR  38
TN1(OsTN8t001161.1)/1-206              1 TNYTKLTTLVLKGLKRVATTQPASSSEIVHEIHSKNSWNKEMRLSTCSTALDSTLKTCSR  60

IR8(OsIR8_08T0127600.1)/1-194         39 FIS-----------------LWLIGNNLHNLELWRLEVASHDLHVVKYLLDLLRLNYEDV  81
MH63(OsMH_08T0238500.1)/1-194         39 FIS-----------------LWLIGNNLHNLELWRLEVASHDLHVVKYLLDLLRLNYEDV  81
O_nivara(ONIVA04G28960.1)/1-201       39 FISPFILFVQSYPFPDIDGQLWLIGNNLHNLELWRLEVASHDLHVVKYLLDLLRLNYEDV  98
O_rufipogon(ORUFI08G12790.1)/1-184    39 FIS-----------------LWLIGNNLHNLELWRLEVASHDLHVVKYLLDLLGLNYEDV  81
TN1(OsTN8t001161.1)/1-206             61 FIS-----------------LWLIGNNLHNLELWRLEVASHDLHVVKYLLDLLRLNYEDV 103

IR8(OsIR8_08T0127600.1)/1-194         82 LPVYIGDDTTDENAFKVLDSGPIYIGDDTTDENAFKEKLQQAHDIFYFCKVHEIHLKDSR 141
MH63(OsMH_08T0238500.1)/1-194         82 LPVYIGDDTTDENAFKVLDSGPIYIGDDTTDENAFKEKLQQAHDIFYFCKVHEIHLKDSR 141
O_nivara(ONIVA04G28960.1)/1-201       99 LPVYIGDDTTDENAFKVLDSGPIYIGDDTTDENAFKEKLQQAHDIFYFCKVHEIHLKDSR 158
O_rufipogon(ORUFI08G12790.1)/1-184    82 LPIYIGDDTTDENAFKVLDSGPIYIGDDTTDENAFKEKLQQAHDIFYFCKVHEIHLKDSR 141
TN1(OsTN8t001161.1)/1-206            104 LPVYIGDDTTDENAFKVLDSGPIYIGDDTTDENAFKEKLQQAHDIFYFCKVHEIHLKDSR 163

IR8(OsIR8_08T0127600.1)/1-194        142 NKGNAVKYMLDRLGLNSEDVLPIYIGDDTTDENAFKVMEFLCLLASWRELSQS------- 194
MH63(OsMH_08T0238500.1)/1-194        142 NKGNAVKYMLDRLGLNSEDVLPIYIGDDTTDENAFKVMEFLCLLASWRELSQS------- 194
O_nivara(ONIVA04G28960.1)/1-201      159 NKGNAVKYMLDRLGLNSEDVLPIYIGDDTTDENAFK-----------------TITKWDG 201
O_rufipogon(ORUFI08G12790.1)/1-184   142 NKGNAVKYMLDRLGLNSEDVLPIYIGDDTTDENAFK-----------------TITKWDG 184
TN1(OsTN8t001161.1)/1-206            164 NKGNAVKYMLDRLGLNSEDVLPIYIGDDTTDENAFK-----------------TITKWDG 206
```

**Figure S10g. OsTN8g001161.1 (probable TPP C) translated CDS alignment.** The TN1 PS sites and their probability of being under PS: Ser25 (99.59%), Ser26 (98.79%), Glu27 (95.75%), and Ile28 (95.74%).

```
IR64(IR64_t00038716-R1)/1-577        1 MASQHLILLLAIFVSLCVAAIGQGNKIVPFNPSCSTTGNYSGDSQYKKNLDQLLSTLATA  60
IR8(OsIR8_12T0187300.1)/1-589        1 MASQHLILLLAIFVSLCVAAIGQGNKIVPFNPSCSTTGNYSGDSQYKKNLDQLLSTLATA  60
TN1(OsTN12t002058.1)/1-568           1 -------------------AIGQGNKIVPFNPSCSTTGNYSGDSQYKKNLDQLLSTLATA  41
MH63(OsMH_12T0345500.1)/1-619        1 MASQHLILLLAIFVSLCVAAIGQGNKIVPFNPSCSTTGNYSGDSQYKKNLDQLLSTLATA  60

IR64(IR64_t00038716-R1)/1-577       61 ATDDGWFNTSSVGTGGDDQVFGLIMCYADRNPTQCKECLAGAPAGITQVCPGSRTVNANY 120
IR8(OsIR8_12T0187300.1)/1-589       61 ATDDGWFNTSSVGTGGDDQVFGLIMCYADRNPTQCKECLAGAPAGITQVCPGSRTVNANY 120
TN1(OsTN12t002058.1)/1-568          42 ATDDGWFNTSSVGTGGDDQVFGLIMCYADRNPTQCKECLAGAPAGITQVCPGSRTVNANY 101
MH63(OsMH_12T0345500.1)/1-619       61 ATDDGWFNTSSVGTGGDDQVFGLIMCYADRNPTQCKECLAGAPAGITQVCPGSRTVNANY 120

IR64(IR64_t00038716-R1)/1-577      121 DACLLRYSDVSFFSVADKTVAFNVYAKSYVENMAAMNETRWQLMSQLAETAGQTKLRLDT 180
IR8(OsIR8_12T0187300.1)/1-589      121 DACLLRYSDVSFFSVADKTVAFNVYAKSYVENMAAMNETRWQLMSQLAETAGQTKLRLDT 180
TN1(OsTN12t002058.1)/1-568         102 DACLLRYSDVSFFSVADKTVAFNVYAKSYVENMAAMNETRWQLMSQLAETAGQTKLRLDT 161
MH63(OsMH_12T0345500.1)/1-619      121 DACLLRYSDVSFFSVADKTVAFNVYAKSYVENMAAMNETRWQLMSQLAETAGQTKLRLDT 180

IR64(IR64_t00038716-R1)/1-577      181 GSTRLGSTSMMYGLAQCTRDLAVSECSTCLSDYIVQLSKIFPNNSGAAIKGYSCYLRYDL 240
IR8(OsIR8_12T0187300.1)/1-589      181 GSTRLGSTSMMYGLAQCTRDLAVSECSTCLSDYIVQLSKIFPNNSGAAIKGYSCYLRYDL 240
TN1(OsTN12t002058.1)/1-568         162 GSTRLGSTSMMYGLAQCTRDLAVSECSTCLSDYIVQLSKIFPNNSWAAIKGYSCYLRYDL 221
MH63(OsMH_12T0345500.1)/1-619      181 GSTRLGSTSMMYGLAQCTRDLAVSECSTCLSDYIVQLSKIFPNNSGAAIKGYSCYLRYDL 240

IR64(IR64_t00038716-R1)/1-577      241 SPFGITLPPSSPVPPPSSTRSTGFVAGLSVAGAVSFMVILGVSIWLLLRRRRKHARLMRE 300
IR8(OsIR8_12T0187300.1)/1-589      241 SPFGITLPPSSPVPPPSSTRSTGFVAGLSVAGAVSFMVILGVSIWLLLRRRRKHARLMRE 300
TN1(OsTN12t002058.1)/1-568         222 SPFGITLPPSSPVPPPSSTRSTGFVAGLSVAGAVSFMVILGVSIWLLLRRRRKHARLMRE 281
MH63(OsMH_12T0345500.1)/1-619      241 SPFGITLPPSSPVPPPSSTRSTGFVAGLSVAGAVSFMVILGVSIWLLLRRRRKHARLMRE 300

IR64(IR64_t00038716-R1)/1-577      301 HQEMEDDFEKGTRPKRFRYDELSVATDFFSDDCKLGEGGF-------------------- 340
IR8(OsIR8_12T0187300.1)/1-589      301 HQEMEDDFEKGTRPKRFRYDELSVATDFFSDDCKLGEGGFGSKQGRKEYESEVRIISRLR 360
TN1(OsTN12t002058.1)/1-568         282 HQEMEDDFEKGTRPKRFRYDELSVATDFFSDDCKLGEG--GSKQGRKEYESEVRIISRLR 339
MH63(OsMH_12T0345500.1)/1-619      301 HQEMEDDFEKGTRPKRFRYDELSVATDFFSDDCKLGEGGFGSKQGRKEYESEVRIISRLR 360

IR64(IR64_t00038716-R1)/1-577      341 ------GSVYKGFLKDLNELIGWCHDGSELLLVYELMPNASLDTHLYNANANVLPWPLR 394
IR8(OsIR8_12T0187300.1)/1-589      361 HRNLVQ--------------LIGWCHDGSELLLVYELMPNASLDTHLYNANANVLPWPLR 406
TN1(OsTN12t002058.1)/1-568         340 HRNLVQ--------------LIGWCHDGSELLLVYELMPNASLDTHLYNANANVLPWPLR 385
MH63(OsMH_12T0345500.1)/1-619      361 HRNLVQ--------------LIGWCHDGSELLLVYELMPNASLDTHLYNANANVLPWPLR 406

IR64(IR64_t00038716-R1)/1-577      395 YEIVLGTGSALLYLHEEWEQCVVHRDIKPSNIMLDTAFNAKLGDFGLARLVDHDR----- 449
IR8(OsIR8_12T0187300.1)/1-589      407 YEIVLGTGSALLYLHEEWEQCVVHRDIKPSNIMLDTAFNAKLGDFGLARLVDHDR----- 461
TN1(OsTN12t002058.1)/1-568         386 YEIVLGTGSALLYLHEEWEQCVVHRDIKPSNIMLDTAFNAKLGDFGLARLVDHDR----- 440
MH63(OsMH_12T0345500.1)/1-619      407 YEIVLGTGSALLYLHEEWEQCVVHRDIKPSNIMLDTAFNAKLGDFGLARLVDHDRAGTMG 466

IR64(IR64_t00038716-R1)/1-577      450 ------------------------------GSHTNADEEEDMIHIAQWVWDLYGNGRIID 479
IR8(OsIR8_12T0187300.1)/1-589      462 ------------------------------GSHTNADEEEDMIHIAQWVWDLYGNGRIID 491
TN1(OsTN12t002058.1)/1-568         441 ------------------------------GSHTNADEEEDMIHIAQWVWDLYGNGRIID 470
MH63(OsMH_12T0345500.1)/1-619      467 YMDPECMITGRANAESDVYSFGVVLLEIAC-----ADEEEDMIHIAQWVWDLYGNGRIID 521

IR64(IR64_t00038716-R1)/1-577      480 AADHRLGGEFNGEEMEAVMVVGLWCAHPDRSLRPTIRQAVSVLQGEVPPPSLPTRMPVAT 539
IR8(OsIR8_12T0187300.1)/1-589      492 AADHRLGGEFNGEEMEAVMVVGLWCAHPDRSLRPTIRQAVSVLQGEVPPPSLPTRMPVAT 551
TN1(OsTN12t002058.1)/1-568         471 AADHRLGGEFNGEEMEAVMVVGLWCAHPDRSLRPTIRQAVSVLRGEVPPPSLPTRMPVAT 530
MH63(OsMH_12T0345500.1)/1-619      522 AADHRLGGEFNGEEMEAVMVVGLWCAHPDRSLRPTIRQAVSVLQGEVPPPSLPTRMPVAT 581

IR64(IR64_t00038716-R1)/1-577      540 FLSPVDAFNYTSSYVTGNTNTSTSTNTTQSSRTTGTVA                       577
IR8(OsIR8_12T0187300.1)/1-589      552 FLSPVDAFNYTSSYVTGNTNTSTSTNTTQSSRTTGTVA                       589
TN1(OsTN12t002058.1)/1-568         531 FLSPVDAFNYTSSYVTGNTNTSTSTNTTQSSRTTGTVA                       568
MH63(OsMH_12T0345500.1)/1-619      582 FLSPVDAFNYTSSYVTGNTNTSTSTNTTQSSRTTGTVA                       619
```

**Figure S10h. OsTN12g002058.1 (L-type lectin-domain containing receptor kinase IX.1-like) translated CDS alignment.** The TN1 PS sites and their probability of being under PS: Trp207 (63.59%) and Arg514 (43.54%).

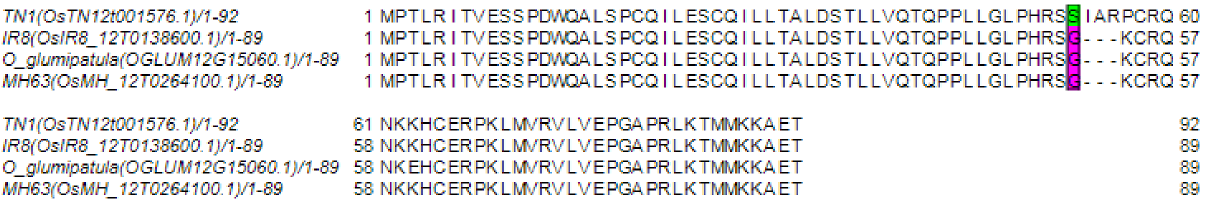


**Figure S10i. OsTN12g001576.1 translated CDS alignment.** The TN1 PS site and the probability of being under PS: Ser53 (85.69%).

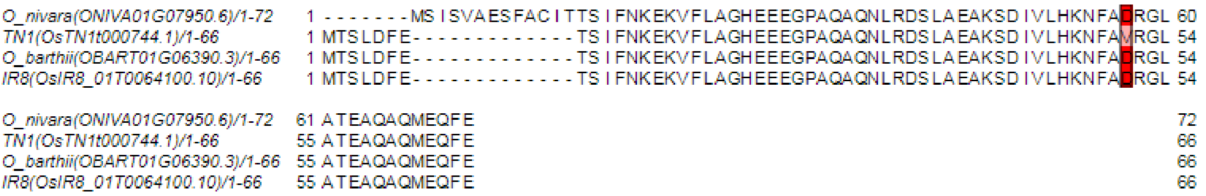


**Figure S10j. OsTN1g000744.1 (KARI, chloroplastic) translated CDS alignment.** The TN1 PS site and the probability of being under PS: Val51 (90.33%).

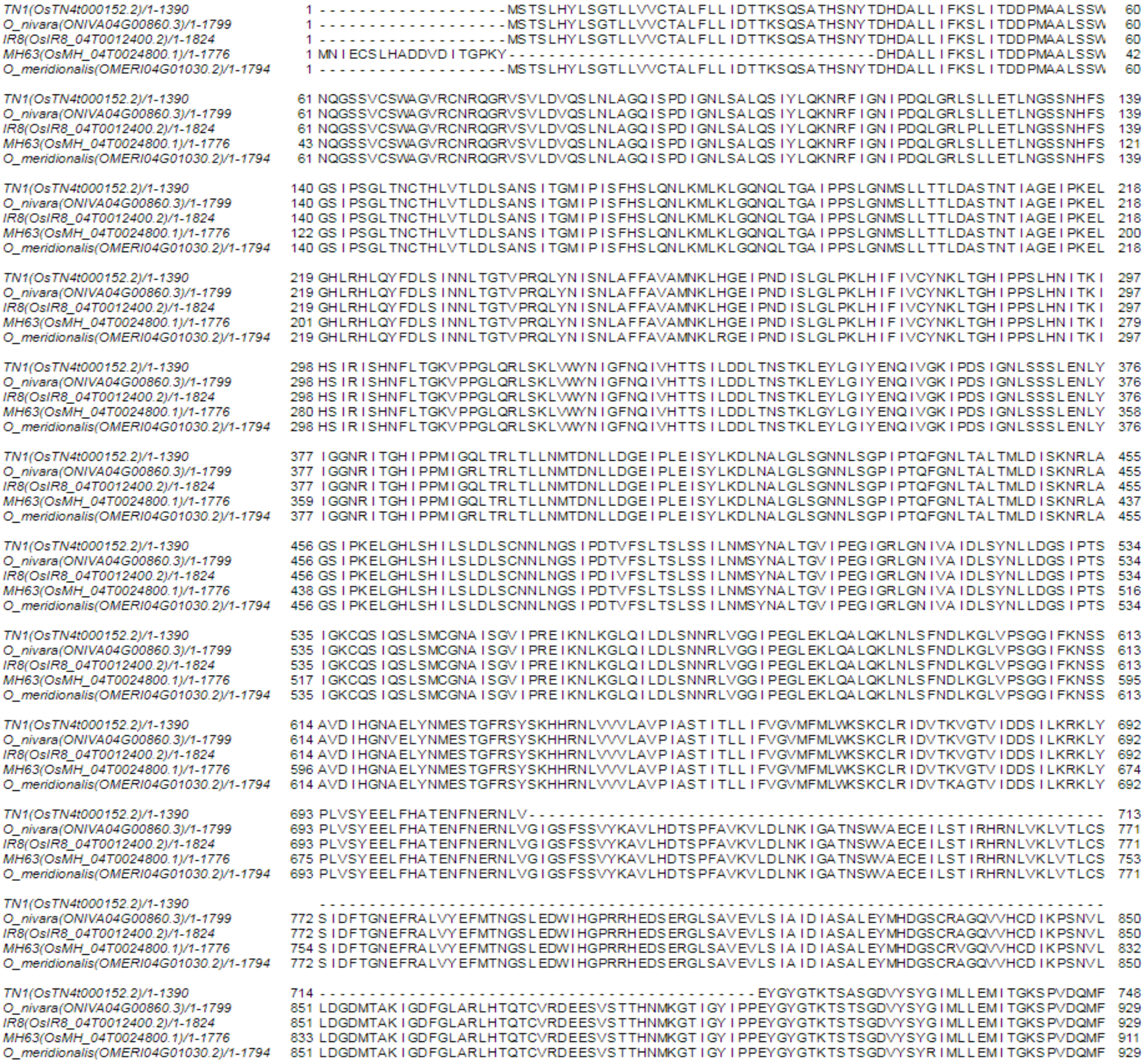

**Figure S10k. OsTN4g000152.2 (probable LRR receptor-like serine/threonine-protein kinase At3g47570).** The TN1 PS sites and their probability of being under PS: Lys1182 (69.84%) and Thr1286 (60.54%).

```
TN1(OsTN4t000152.2)/1-1390                749 EGEMNLEKWVRVSIPHQADEVVDKRFMITGSEESSADGQQQQQVDTVDSKLLLETLLVPMVDVALCCVRESPGSRISMH 827
O_nivara(ONIVA04G00860.3)/1-1799          930 EGEMNLEKWVRVSIPHQADEVVDKRFLITGSEESSADGQQQQQVDTVDSKLLLETLLVPMVDVALCCVRESPGSRISMH 1008
IR8(OsIR8_04T0012400.2)/1-1824            930 EGEMNLEKWVRVSIPHQADEVVDKRFLITGSEESSADGQQQQQVDTVDSKLLLETLLVPMVDVALCCVRESPGSRISMH 1008
MH63(OsMH_04T0024800.1)/1-1776            912 EGEMNLEKWVRVSIPHQADEVVDKRFMITGSEESSADGQQQQQVDTVDSKLLLETLLVPMVDVALCCVRESPGSRISMH 990
O_meridionalis(OMERI04G01030.2)/1-1794    930 EGEMNLEKWVRVSIPHQADEVVDKRFMITGSEESSADGQQQQQVDTVDSKLLLETLLVPMVDVALCCVRESPDSRISMH 1008

TN1(OsTN4t000152.2)/1-1390                828 DALSRLKRINEKFLKSLA------------------------------VCRSCSDGERHALLRRIQPLIGPEFSSNGRL 876
O_nivara(ONIVA04G00860.3)/1-1799         1009 DALSRLKRINEKFLKSLA------------------------------VCRSCSDGERHALLRRIQPLIGPEFSSNGRL 1057
IR8(OsIR8_04T0012400.2)/1-1824           1009 DALSRLKRINEKFLKSLALGLAPSGYVDTNTVAVVLPPANGDKTFHHFVCRSCSDGERHALLRRIQPLIGPEFSSNGRL 1087
MH63(OsMH_04T0024800.1)/1-1776            991 DALSRLKRINEKFLKSLA------------------------------VCRSCSDGERHALLRRIQPLIGPEFSSNGRL 1039
O_meridionalis(OMERI04G01030.2)/1-1794   1009 DALSRLKRINEKFLKSLA------------------------------VCRSCSDGERHALLRRIQPLIGPEFSSNGRL 1057

TN1(OsTN4t000152.2)/1-1390                877 DWHEAVDCCRWEGVTCSVAGRRREAAGGRRVVSLSLPGVGIAGAVDAAVLAPFTALEKLDLSGNQITSFSAANRSDMVV 955
O_nivara(ONIVA04G00860.3)/1-1799         1058 DWHEAVDCCRWEGVTCSVAGRRREAAGGRRVVSLSLPGVGIAGAVDAAVLAPFTALEKLDLSGNQITSFSAANRSDMVV 1136
IR8(OsIR8_04T0012400.2)/1-1824           1088 DWHEAVDCCRWEGVTCSVAGRRREAAGGRRVVSLSLPGVGIAGAVDAAVLAPFTALEKLDLSGNQITSFSAANRSDMVV 1166
MH63(OsMH_04T0024800.1)/1-1776           1040 DWHEAVDCCRWEGVTCSVAGRRREAAGGRRVVSLSLPGVGIAGAVDAAVLAPFTALEKLDLSGNQITSFSAANRSDMVV 1118
O_meridionalis(OMERI04G01030.2)/1-1794   1058 DWHEAVDCCRWEGVTCSVAGRRREAAGGRRVVSLSLPGVGIAGAVDAAVLAPFTALEKLDLSGNQITSFSAANRSDMVV 1136

TN1(OsTN4t000152.2)/1-1390                956 GAVLNNLTALTELHLAGNEITTTGWISNLTSLQVIDMSSNKLHELN------------------------GVINPCLGK 1010
O_nivara(ONIVA04G00860.3)/1-1799         1137 GAVLNNLTALTELHLAGNEITTTGWISNLTSLQVIDMSSNKLHELNAPIYTGICGLHQLKYLSVGFNMIQGVINPCLGK 1215
IR8(OsIR8_04T0012400.2)/1-1824           1167 GAVLNNLTALTELHLAGNEITTTGWISNLTSLQVIDMSSNKLHELN-----GICGLHQLKYLSVGFNMIQGVINPCLGK 1240
MH63(OsMH_04T0024800.1)/1-1776           1119 GAVLNNLTALTELHLAGNEITTTGWISNLTSLQVIDMSSNKLHELN-----GICGLHQLKYLSVGFNMIQGVINPCLGK 1192
O_meridionalis(OMERI04G01030.2)/1-1794   1137 GAVLNNLTALTELHLAGNEITTTGWISNLTSLQVIDMSSNKLHELN-----GICGLHQLKYLSVGFNMIQGVINPCLGK 1210

TN1(OsTN4t000152.2)/1-1390               1011 LQHLVYLDMGSNFLTGEIGQYLLSNLTQVEEVHLGDNNLTGTFDFSSLANNSELHSIVLSNNCKLEIETELVRWTPLFQ 1089
O_nivara(ONIVA04G00860.3)/1-1799         1216 LQHLVYLDMGSNFLTGEIGQNLLSNLTRVEQVHLGDNNLTGTFDFSSLANNSELHSIVLSNNYKLEIETELVRWTPLFQ 1294
IR8(OsIR8_04T0012400.2)/1-1824           1241 LQHLVYLDMGSNFLTGEIGQNLLSNLTRVEQVHLGDNNLTGTFDFSSLANNSELHSIVLSNNYKLEIETELVRWTPLFQ 1319
MH63(OsMH_04T0024800.1)/1-1776           1193 LQHLVYLDMGSNFLTGEIGQNLLSNLTRVEQVHLGDNNLTGTFDFSSLANNSELHSIVLLNNYKLEIETELVRWTPLFQ 1271
O_meridionalis(OMERI04G01030.2)/1-1794   1211 LQHLVYLDMGSNFLTGEIGQNLLSNLTRVEQVHLGDNNLTGTFDFSSLANNSELHSIVLSNNYKLEIETELVRWTPLFQ 1289

TN1(OsTN4t000152.2)/1-1390               1090 LEYLNLSNTIVNKRSNGIIPTFLSAQVSLSGIDLSICSLQGRIPSWMLLYNVSLGFLLLRGNSMDFLDTGNLGANVTSS 1168
O_nivara(ONIVA04G00860.3)/1-1799         1295 LEYLNLSNSIVNKRSNGIIPTFLSAQVSLSGIDLSICSLQGRIPSSMLLYNVSLGFLLLRGNSMDFLDTGNLGANVTSS 1373
IR8(OsIR8_04T0012400.2)/1-1824           1320 LEYLNLSNSIVNKRSNGIIPTFLSAQVSLSGIDLSICSLQGRIPSWMLLYNVSLGFLLLRGNSMDFLDTGNLGANVTSS 1398
MH63(OsMH_04T0024800.1)/1-1776           1272 LEYLNLSNSIVNKRSNGIIPTFLSAQVSLSGIDLSICSLQGRIPSSMLLYNVSLGFLLLRGNSMDFLDTGNLGANVTSS 1350
O_meridionalis(OMERI04G01030.2)/1-1794   1290 LEYLNLSNSIVNKRSNGIIPTFLSAQVSLSGIDLSICSLQGRIPSWMLLYNVSLGFLLLRGNSMDFLDTGNLGANVTSS 1368

TN1(OsTN4t000152.2)/1-1390               1169 MKVLDLSDNRISMKMPYNFGSLFPYLKYLDMSSNMLNGGVPSLAEAASSLQVLDLSFNMLNGEISPELIGNASILTSLL 1247
O_nivara(ONIVA04G00860.3)/1-1799         1374 MEVLDLSNNMISMPMPYNLGSLFPYLKYLDMSSNMLHGGVPSLAEAVSSLQVLDLSFNRLDGEISPEFIGNASILTSLL 1452
IR8(OsIR8_04T0012400.2)/1-1824           1399 MEVLDLSNNMISMPMPYNLGSLFPYLKYLDMSSNMLHGGVPSLAEAVSSLQVLDLSFNRLDGEISPEFIGNASILTSLL 1477
MH63(OsMH_04T0024800.1)/1-1776           1351 MEVLDLSNNMISMPMPYNLGSLFPYLKYLDMSSNMLHGGVPSLAEAVSSLQVLDLSFNRLDGEISPEFIGNASILTSLL 1429
O_meridionalis(OMERI04G01030.2)/1-1794   1369 MEVLDLSNNMISMPMPYNLGSLFPYLKYLDMSSNMLHGGVPSLAEAVSSLQVLDLSFNRLDGEISPEFIGNASILTSLL 1447

TN1(OsTN4t000152.2)/1-1390               1248 LSHNDLTGPMPPFHWSLGQLTHLNVKNNQLSGRLPPLLTNCTNLENLNVRNNRLSGVIPVGLLSFEKLGALLLGGNQFH 1326
O_nivara(ONIVA04G00860.3)/1-1799         1453 LSHNDLTGPMPPFHWIPGQLIHLSIENNQLSGGLPPLLMNCTNLENLNVRNNRLSGVIPVGLLNFEKLGALLLGGNQFH 1531
IR8(OsIR8_04T0012400.2)/1-1824           1478 LSHNDLTGPMPPFHWIPGQLIHLSIENNQLSGGLPPLLMNCTNLENLNVRNNRLSGVIPVGLLNFEKLGALLLGGNQFH 1556
MH63(OsMH_04T0024800.1)/1-1776           1430 LSHNDLTGPMPPFHWIPGQLIHLSIENNQRSGGLPPLLMNCTNLENLNVRNNRLSGVIPVGLLNFEKLGALLLGGNQFH 1508
O_meridionalis(OMERI04G01030.2)/1-1794   1448 LSHNDLTGPMPPFHWIPGQLIHLSVENNQLSGGLPPLLMNCTNLENLNVRNNRLSGVITVGLLNFEKLGALLLGGNQFH 1526

TN1(OsTN4t000152.2)/1-1390               1327 GVIPWDICLNNYIHFIDLSNNWFSGEIPGCLYS-----LPMYYEDDPFGNITQRRQTYVEFTTKGESLT---------- 1390
O_nivara(ONIVA04G00860.3)/1-1799         1532 GVIPWDICLNNNLHFIDLSNNRFSGEIPGCLYSVFWSELPMYYEDDPFGNITQRRQTYVEFTTKGESLTYMGMPLELMT 1610
IR8(OsIR8_04T0012400.2)/1-1824           1557 GVIPWDICLNNNLHFIDLSNNRFSGEIPGCLYSVFWSELPMYYEDDPFGNITQRRQTYVEFTTKGESLTYMGMPLELMT 1635
MH63(OsMH_04T0024800.1)/1-1776           1509 GVIPWDICLNNNLHFIDLSNNRFSGEIPGCLYSVFWSELPMYYEDDPFGNITQRRQTYVEFTTKGESLTYMGMPLELMT 1587
O_meridionalis(OMERI04G01030.2)/1-1794   1527 GVIPWDICLNNNLHFIDLSNNQFSGEIPGCLSSVFWSELPMYYEDDPFGNITQRRQTYVEFTTKGESLTYMGMPLELMT 1605

TN1(OsTN4t000152.2)/1-1390                    -------------------------------------------------------------------------------
O_nivara(ONIVA04G00860.3)/1-1799         1611 GIDLSMNRLSGTIPSPIGFLRQLKSLNLSHNKLVGSIPDTFMYLLEMESMDLSHNHLNGSVPVELANLSFLSFFSVAYN 1689
IR8(OsIR8_04T0012400.2)/1-1824           1636 GIDLSMNRLSGTIPSPIGFLRQLKSLNLSHNKLVGSIPDTFMYLLEMESMDLSHNHLNGSVPVELANLSFLSFFSVAYN 1714
MH63(OsMH_04T0024800.1)/1-1776           1588 GIDLSMNRLSGTIPSPIGFLRQLKSLNLSHNKLVGSIPDTFMYLLEMESMDLSHNHLNGSVPVELANLSFLSFFSVAYN 1666
O_meridionalis(OMERI04G01030.2)/1-1794   1606 GIDLSMNRLSGTIPSPVGFLRQLKSLNLSHNKLVGSIPDTFMYLLEMESMDLSHNHLNGSIPVELANLSFLSFFSVAYN 1684

TN1(OsTN4t000152.2)/1-1390                    -------------------------------------------------------------------------------
O_nivara(ONIVA04G00860.3)/1-1799         1690 NLSGEIPFESQLCTLNGTAFEGNENLCGKIVDKICLMNSNHSHDSDDEMHQLLSTDTMDTPLIYWSFVAGSFAIGFWGI 1768
IR8(OsIR8_04T0012400.2)/1-1824           1715 NLSGEIPFESQLCTLNGTAFEGNENLCGEIVDKICLMNSNHSHDSDDEMHQLLSTDTMDTPLIYWSFVAGSFAIGFWGI 1793
MH63(OsMH_04T0024800.1)/1-1776           1667 NLSGEIPFESQLCTLNGTAFEGNENLCGKIVDKICLMNSNHSHDSDDEMHQLLSTDTMDTPLIYWSFVAGSFAIGFWGI 1745
O_meridionalis(OMERI04G01030.2)/1-1794   1685 NLSGEIPFESQLCTLNGTAFEGNENLRGDIFDKICLMNSNHSHDSDDEMHQLLSTDTMDTPLIYWFFVAGSFAIGFWGI 1763

TN1(OsTN4t000152.2)/1-1390                    -------------------------------
O_nivara(ONIVA04G00860.3)/1-1799         1769 IALLIWNTTFRSRLCSFMDGCMSKMGWILVP                                                 1799
IR8(OsIR8_04T0012400.2)/1-1824           1794 IALLIWNTTFRSRLCSFMDGCMSKMGWILVP                                                 1824
MH63(OsMH_04T0024800.1)/1-1776           1746 IALLIWNTTFRSRLCSFMDGCMSKMGWILVP                                                 1776
O_meridionalis(OMERI04G01030.2)/1-1794   1764 IALLIWNTTFRSRLCSFMDGCMSKMGWILVP                                                 1794
```

**Figure S10k continued.**

#### I. SUPPLEMENTARY TABLES

**Table S1. Number of Tetep NLRs that are found in TN1, MH63, R498 and Nipponbare.** The sum of the NLRS with orthologues and with blastp hits ≥ 50%, (429 for TN1, 423 for MH63, 436 for R498, 438 for Nipponbare), did not differ much.

| Tetep (455 NLRs) | | TN1 | MH63 | R498 | Nipponbare |
|---|---|---|---|---|---|
| NLRs with orthologues(defined by OrthoFinder) | | 322 | 335 | 332 | 360 |
| NLRs without orthologues | Blastp Identity ≥ 50% | 107 | 88 | 104 | 78 |
| | Blastp Identity < 50% | 26 | 32 | 19 | 17 |

**Table S2: Proportion of R genes that have been tested either in TP309 or Shin2.** The numbers on this table were used in the Chi-square computation in R.

| | Tested | Resistant (either TP309 or Shin2) |
|---|---|---|
| **Tetep NBS** | 219 | 90 |
| **Orthologs in TN1(defined by OrthoFinder)** | 170 | 69 |

**Table S3. Gene Ontology (GO) terms of the TN1 genes under positive selection.** The GOs on this table were used as input in REVIGO.

| Gene name | protein name | GO id | meaning |
|---|---|---|---|
| OsTN5g000040 | hypothetical protein | - | - |
| OsTN5g002486 | - | - | - |
| OsTN2g002903 | PLATZ transcription factor family protein | - | - |
| OsTN5g001087 | GA 3beta-hydroxylase | Biological Process | |
| | | GO:0009416 | response to light stimulus |
| | | GO:0009686 | gibberellin biosynthetic process |
| | | GO:0009826 | unidimensional cell growth |
| | | GO:0009908 | flower development |
| | | GO:0055114 | oxidation-reduction process |
| | | Molecular Function | |
| | | GO:0016707 | gibberellin 3-beta-dioxygenase activity |
| | | GO:0045544 | gibberellin 20-oxidase activity |
| OsTN1g003572 | armadillo/beta-catenin repeat protein-like | - | - |
| OsTN1g003413 | transmembrane protein 56 isoform X1 | Biological Process | |
| | | GO:0055085 | transmembrane transport |
| | | Cellular Component | |
| | | GO:0005886 | plasma membrane |
| | | GO:0016021 | integral component of membrane |
| | | Molecular Function | |
| | | GO:0022857 | transmembrane transporter activity |
| OsTN8g001161 | probable trehalose-phosphate phosphatase C | Biological Process | |
| | | GO:0005992 | trehalose biosynthetic process |
| | | GO:0016311 | dephosphorylation |
| | | Molecular Function | |
| | | GO:0004805 | trehalose-phosphatase activity |
| OsTN12g002058 | L-type lectin-domain containing receptor kinase IX.1-like | Biological Process | |
| | | GO:0002229 | defense response to oomycetes |
| | | GO:0006468 | protein phosphorylation |

| | | | |
|---|---|---|---|
| | | GO:0042742 | defense response to bacterium |
| | | Cellular Component | |
| | | GO:0016021 | integral component of membrane |
| | | GO:0005886 | plasma membrane |
| | | Molecular Function | |
| | | GO:0004675 | transmembrane receptor protein serine/threonine kinase activity |
| | | GO:0005524 | ATP binding |
| OsTN12g001576 | - | - | - |
| OsTN1g000744 | Ketol-acid reductoisomerase, chloroplastic | Biological Process | |
| | | GO:0009097 | isoleucine biosynthetic process |
| | | GO:0009099 | valine biosynthetic process |
| | | GO:0055114 | oxidation-reduction process |
| | | Cellular Component | |
| | | GO:0005739 | mitochondrion |
| | | GO:0009507 | chloroplast |
| | | Molecular Function | |
| | | GO:0046872 | metal ion binding |
| | | GO:0004455 | ketol-acid reductoisomerase activity |
| OsTN4g000152 | probable LRR receptor-like serine/threonine-protein kinase At3g47570 | Biological Process | |
| | | GO:0006468 | protein phosphorylation |
| | | GO:0009755 | hormone-mediated signaling pathway |
| | | Cellular Component | |
| | | GO:0005886 | plasma membrane |
| | | GO:0016021 | integral component of membrane |
| | | Molecular Function | |
| | | GO:0004674 | protein serine/threonine kinase activity |
| | | GO:0005515 | protein binding |
| | | GO:0005524 | ATP binding |

**Table S4. Statistics of coding sequences used in the PosiGene run and their source link.** With the exception of TN1, IR64 and MH63, all the coding sequences of the input species were downloaded from the Gramene ftp website. Because the ftp link can change without prior notice, users wanting to download the same data should refer to release 62 and oge release 3, if ever the links had changed.

| organism | Version/ Date downloaded or extracted | # CDS | Total length of coding sequences | source link |
|---|---|---|---|---|
| TN1 | April 26, 2020 | 37,952 | 45,684,501 | - |
| IR8 | May 10, 2020 | 56,823 | 76,985,769 | http://ftp.gramene.org/oge/release-3/fasta/oryza_indicair8/dna/ |
| IR64 | May 10, 2020 | 41,458 | 45,637,530 | https://rootomics.dna.affrc.go.jp/en/research/IR64 |
| MH63 | MH63RS2 | 83,258 | 109,797,326 | http://rice.hzau.edu.cn/cgi-bin/rice_rs2/download_ext |
| *Oryza nivara* | v1.0 | 48,360 | 61,006,575 | http://ftp.gramene.org/archives/PAST_RELEASES/release-62/fasta/oryza_nivara/cds/ |
| *Oryza rufipogon* | May 10, 2020 | 50,219 | 59,474,618 | http://ftp.gramene.org/oge/release-3/fasta/oryza_rufipogon/dna/ |
| 9311 | ASM465v1 | 40,745 | 45,520,686 | http://ftp.gramene.org/archives/PAST_RELEASES/release-62/fasta/oryza_indica/cds/ |
| Nipponbare | March 24, 2020 | 42,313 | 42,136,737 | https://rapdb.dna.affrc.go.jp/download/irgsp1.html |
| *Oryza barthii* | v1 | 41,595 | 50,568,141 | http://ftp.gramene.org/archives/PAST_RELEASES/release-62/fasta/oryza_barthii/cds/ |
| *Oryza brachyantha* | v1.4b | 32,037 | 34,004,031 | http://ftp.gramene.org/archives/PAST_RELEASES/release-62/fasta/oryza_brachyantha/cds/ |
| *Oryza glaberrima* | V1 | 33,164 | 36,293,721 | http://ftp.gramene.org/archives/PAST_RELEASES/release-62/fasta/oryza_glaberrima/cds/ |
| *Oryza glumipatula* | v1.5 | 46,893 | 58,081,746 | http://ftp.gramene.org/archives/PAST_RELEASES/release-62/fasta/oryza_glumipatula/cds/ |
| *Oryza punctata* | v1.2 | 41,060 | 53,482,920 | http://ftp.gramene.org/archives/PAST_RELEASES/release-62/fasta/oryza_punctata/cds/ |
| *Oryza meridionalis* | v1.3 | 43,455 | 55,214,595 | http://ftp.gramene.org/archives/PAST_RELEASES/release-62/fasta/oryza_meridionalis/cds/ |
| *Oryza longistaminata* | v1.0 | 31,686 | 34,731,219 | http://ftp.gramene.org/archives/PAST_RELEASES/release-62/fasta/oryza_longistaminata/cds/ |
| *Brachypodium distachyon* | v3.0 | 52,972 | 66,747,974 | http://ftp.gramene.org/archives/PAST_RELEASES/release-62/fasta/brachypodium_distachyon/cds/ |

| | | | | |
|---|---|---|---|---|
| *Eragrostis tef* | ASM97063v1 | 41,555 | 48,850,019 | http://ftp.gramene.org/archives/PAST_RELEASES/release-62/fasta/eragrostis_tef/cds/ |
| *Leersia perrieri* | V1.4 | 38,960 | 52,135,503 | http://ftp.gramene.org/archives/PAST_RELEASES/release-62/fasta/leersia_perrieri/cds/ |
| *Panicum hallii fil2* | v3.1 | 44,192 | 54,354,546 | http://ftp.gramene.org/archives/PAST_RELEASES/release-62/fasta/panicum_hallii_fil2/cds/ |
| *Panicum hallii hal2* | v2.1 | 42,523 | 51,996,270 | http://ftp.gramene.org/archives/PAST_RELEASES/release-62/fasta/panicum_hallii_hal2/cds/ |
| *Setaria italica* | v2.0 | 41,023 | 45,897,567 | http://ftp.gramene.org/archives/PAST_RELEASES/release-62/fasta/setaria_italica/cds/ |
| *Sorghum bicolor* | NCBIv3 | 47,110 | 57,859,353 | http://ftp.gramene.org/archives/PAST_RELEASES/release-62/fasta/sorghum_bicolor/cds/ |
| *Triticum aestivum* | IWGSC | 133,346 | 177,672,347 | http://ftp.gramene.org/archives/PAST_RELEASES/release-62/fasta/triticum_aestivum/cds/ |
| *Zea mays* | v4 | 131,585 | 190,385,016 | http://ftp.gramene.org/archives/PAST_RELEASES/release-62/fasta/zea_mays/cds/ |

**Dataset S1:** Predicted R genes in TN1, their Pfam domains, NLR-Parser result and R gene classification

**Dataset S2:** Results of finding Tetep NLRs in TN1

**Dataset S3:** Blastp and tblastn hits of the cloned R genes to the TN1 and Tetep genome

**Dataset S4:** Haplotype and variety order of Pi54 and Pi-ta from SNP-Seek

**Dataset S5:** SNP effect results from SNP-Seek
